# IL-22 promotes acute kidney injury through activation of the DNA damage response and cell death in proximal tubule cells

**DOI:** 10.1101/2023.06.08.544134

**Authors:** Kensei Taguchi, Sho Sugahara, Bertha C. Elias, Navjot Pabla, Guillaume Canaud, Craig R. Brooks

## Abstract

Acute kidney injury (AKI) affects over 13 million people world-wide annually and is associated with a fourfold increase in mortality. Our lab and others have shown that DNA damage response (DDR) governs the outcome of AKI in a bimodal manner. Activation of DDR sensor kinases protects against AKI, while hyperactivation of DDR effector proteins, such as p53, induces to cell death and worsens AKI. The factors that trigger the switch from pro-reparative to pro-cell death DDR remain to be resolved. Here we investigate the role of interleukin 22 (IL-22), an IL-10 family member whose receptor (IL-22RA1) is expressed on proximal tubule cells (PTCs), in DDR activation and AKI. Using cisplatin and aristolochic acid (AA) induced nephropathy as models of DNA damage, we identify PTCs as a novel source of urinary IL-22, making PTCs the only epithelial cells known to secret IL-22, to our knowledge. Functionally, IL-22 binding its receptor (IL-22RA1) on PTCs amplifies the DDR. Treating primary PTCs with IL-22 alone induces rapid activation of the DDR *in vitro*. The combination of IL-22 + cisplatin or AA treatment on primary PTCs induces cell death, while the same dose of cisplatin or AA alone does not. Global deletion of IL-22 protects against cisplatin or AA induced AKI. IL-22 deletion reduces expression of components of the DDR and inhibits PTC cell death. To confirm PTC IL-22 signaling contributes to AKI, we knocked out IL-22RA1 in renal epithelial cells by crossing IL-22RA1floxed mice with Six2-Cre mice. IL-22RA1 KO reduced DDR activation, cell death, and kidney injury. These data demonstrate that IL-22 promotes DDR activation in PTCs, switching pro-recovery DDR responses to a pro-cell death response and worsening AKI. Targeting IL-22 represents a novel therapeutic approach to prevent the negative consequences of the DDR activation while not interfering with the processes necessary for repair of damaged DNA.

**Translational statement:** Acute kidney injury, which affects 10-20% of hospitalized patients, is associated with a fourfold increase in mortality and predisposes patients to chronic kidney disease. In the present study, we identify interleukin 22 as a cofactor which worsens acute kidney injury. Interleukin 22 activates the DNA damage response, which in combination with nephrotoxic drugs amplifies the injury response in kidney epithelial cells and increases cell death. Deletion of interleukin 22 from mice or its receptor from mouse kidneys ameliorates cisplatin induced nephropathy. These findings may help clarify the molecular mechanisms of DNA damage induced kidney injury and identify interventions that can help treat acute kidney injury.

## Introduction

Acute kidney injury (AKI) occurs in 21.6% of hospitalized adults worldwide^1^. DNA damage is one of the major manifestations of AKI^2, 3^. Cells respond to DNA damage by activating a sensitive and complex DNA damage response (DDR) pathway, which detects lesions in damaged chromatin, delays cell-cycle progression, and repairs the lesions, or in the case of severe injury promotes cell death. Ataxia telangiectasia mutated (ATM) and ataxia telangiectasia and Rad3-related protein (ATR) are major DDR sensor kinases that sense DNA damage and transmit DDR signals to downstream effector proteins, such as p53. ATM can stabilize p53 through inhibition of the degrader Mouse double minute 2 homolog (MDM2), while both ATM and ATR can activate p53 through phosphorylation. Activation of p53 ultimately regulates cell fate by inducing cell cycle regulators, such as p21, to promote cell cycle arrest or pro-apoptotic genes, such as P53 up-regulated modulator of apoptosis (PUMA) or BCL2 Associated X (BAX), to induce apoptosis^4^. Our lab, and others, have shown that inhibition of p53 is protective against multiple forms of AKI, while inhibition of ATM or ATR dramatically worsens AKI^5–7^. Thus, balanced DDR signaling is necessary for recovery from AKI and prevention of excessive cell death in AKI.

Interleukin-22 (IL-22) is a member of the IL-10–related family of cytokines^8, 9^. IL-22 is predominantly expressed by T cell subsets, such as CD4+, Th17, Th22, T helper, or group 3 innate lymphoid cells (ILC3) [20]. IL-22 receptor complex consists of a ligand-binding chain, the IL-22RA1 and a signal-transducing chain, the IL-10R2. In contrast to ubiquitous expression of IL-10R2^10^, the expression of IL-22RA1 is restricted to epithelial cells, thus, allowing immune-to-epithelial signaling. The binding of IL-22 to IL-22 receptor leads to activation of the transcriptional factor signal transducer and activator 3 (STAT3) via phosphorylation^11, 12^. The primary biological function of IL-22 is regulation of epithelial anti-bacterial responses, through strengthening epithelial barrier integrity^13, 14^. In this way, IL-22 is thought to provide protection against acute injuries, such as ischemia, acetaminophen, or alcohol induced liver injuries by accelerating wound healing^15–17^. On the other hand, IL-22 can induce cell death in intestinal stem cells or erythroid precursors^18, 19^ and plays a pathological role in irritable bowel disease, Crohn’s disease, dermatitis, and lupus nephritis^20–22^. In the clinic, IL-22 inhibition has been shown to be safe and effective at reducing severe atopic dermatitis^23^. Thus, IL-22 signaling can be protective or pathological, depending on the context.

The role of IL-22 In the kidney remains controversial. Earlier studies using high concentrations of IL-22 or IL-22-Fc found IL-22 to be protective against diabetic nephropathy, acetaminophen, and ischemia induced kidney injury^24, 25^, whereas more recent studies have shown endogenous levels of IL-22 or untagged IL-22 worsens lupus nephritis, IgA nephropathy, or hypertensive kidney injury^26, 27^. Previous animal studies, however, have largely relied on systemic overexpression, injection, or global knockout models which modulate IL-22 signaling on the systemic level. Thus, the kidney specific role of IL-22 signaling has yet to be resolved. Also, despite the fact that kidney IL-22RA1 is restricted to the apical surface of PTCs, urinary IL-22 concentration is not monitored in most studies. In the current study, we aim to determine the kidney specific role of IL-22 in nephrotoxin induced AKI.

## Methods

### Animals

Mouse colonies were derived from commercial sources and maintained on a 12-h light/ dark cycle at room temperature with free access to food and water under protocols approved by the institutional animal care and use committee at Vanderbilt University Medical Center. *IL-22RA1flox/flox* (*IL-22RA1fl/fl*; B6.Cg-Il22ra1tm1.1Koll/J) and Six2-TGCtg mice (Six2 Cre; STOCK Tg(Six2-EGFP/Cre)1Amc/J) were purchased from the Jackson laboratory and crossed to create kidney tubule-specific IL-22RA1 knockout mice (*IL-22RA1*^ΔTub^; *IL-22RA1fl/fl; Six2Cre*). *IL-22RA1fl/fl* was utilized to the present study as control. C57BL/6-Il22tm1.1(iCre)Stck/J (IL-22Cre) were purchased from the Jackson laboratory. Exon1 of the Il22 gene was replaced with a codon optimized Cre Recombinase (iCre) which abolishes expression of IL-22 globally in homozygotes. Floxed-stop tdTomato mice (B6.Cg-Gt(ROSA)26Sor^tm14(CAG-tdTomato)Hze^/J) were crossed with IL-22Cre to generate an IL-22 reporter line, IL-22Cre:tdTomato. Experimental mice age 8-12 weeks were divided into control and cisplatin or AA groups at random. Male mice were chosen for the study because we and others have observed a reduced response to both cisplatin and AA induced kidney injury^28, 29^. For cisplatin-induced acute kidney injury, mice were administered cisplatin (Sigma cat#4394) dissolved in sterile saline (KD medical, cat# RGC-3290) at 30 mg/kg in a single intraperitoneal injection. The same amount of sterile saline was injected as control. Blood was taken and body weight measured every 24 hours. For cisplatin studies, wild-type mice were found to reach human endpoints, 20% weight loss Inability reach food/water, hunched posture, and/or abnormal mentation, by day 4-5. For this reason, most mice were sacrificed on day 3 after administration of cisplatin and kidneys, blood, and urine were collected. Aristolochic acid nephropathy (AAN), a model of AKI and subsequent CKD transition, was induced by an intraperitoneal injection of three doses of AA (5 mg/kg body weight, Sigma, cat# A5512) dissolved in sterile saline. The control mice were injected the same amount of saline as control. Blood was collected and body weight measured on day 3, day 7, and every seven days after that until day 42. Mice were sacrificed and kidneys were isolated on day 7 as acute phase and day 42 as chronic phase.

### RNA Scope

RNAScope fluorescent *in situ* hybridization was performed according to the manufacturer’s protocol (ACDBio, RNAscope Multiplex Fluorescent Assay v2). Briefly, 4-6 µm paraffin-embedded kidney sections from patient biopsies or mouse kidneys were deparaffinized and incubated with H2O2 to inhibit endogenous peroxidase activity. RNA was retrieved followed by incubation with proteinase to enable to access to target RNA. Then, the sections were hybridized with *Il-22* probes for 2 hours at room temperature, followed by incubation with appropriate fluorescent probes to visualize the target probe RNA. The images were obtained with a Zeiss LSM 710 confocal microscope and RNA dots were counted by ImageJ (NIH).

### Cytokine staining with Brefeldin A treatment

To identify the source of secreted cytokines, cytokine secretion was inhibited using brefeldin A (BFA) injection and kidneys were stained for cytokines using immunofluorescence staining as previously described^30^. Briefly, brefeldin A was dissolved with dimethyl sulfoxide at a concentration of 10 mg/mL as a stock solution and further diluted with sterile PBS at the final concentration of 1.25 mg/mL immediately before use. 200 μL of a 1.25 mg/mL BFA solution was injected into mice through the tail vein 6 hours prior to sacrifice. Kidneys were harvested and processed for immunofluorescence staining as described above.

### Human samples

Deidentified human biopsies were obtained from 3 patients with cisplatin nephropathy as part of standard clinical practice (Supplemental Table 1). Deidentified human samples from 3 patients with CKD undergoing partial or radical nephrectomy procedure for urological indication. Interstitial fibrosis was evaluated by a trained senior pathologist and the tissue with >10% interstitial fibrosis was categorized as fibrotic kidney (Supplemental Table 2). Kidney biopsies were obtained as part of standard clinical practice. IL-22 staining was performed using RNAScope as described above.

### Study approvals

All animal experiments were approved by the Institutional Animal Care and Use Committee of Vanderbilt University Medical Center. Human samples were obtained under protocols approved by the Necker hospital Institutional Review Board and Boston University Medical Center and Brigham and Women’s Hospital. All necessary patient/participant written informed consent was obtained.

### Statistics

All data were presented as means ±standard deviation (SD). One-way ANOVA and subsequent Tukey’s post hoc test was used to determine statistical differences among over three groups. When two groups were compared, the unpaired two-tailed t-test was conducted. Survival curves were derived using the Kaplan-Meier method and compared using a log-rank test. Statistical analysis was performed using GraphPad Prism 7 software (GraphPad Software, San Diego, CA). For all comparison, p value < 0.05 was considered to indicate statistical significance.

## Results

### PTCs are a novel source of IL-22

To investigate the role of IL-22 in DNA damage induced kidney injury, we stained patient biopsies from patients with cisplatin induced nephropathy for IL-22 RNA using RNAScope (patient characteristics in Supplemental Tables 1 and 2). We were surprised to find that kidney tubule cells express IL-22 RNA (Figure 1A). To confirm these results, we treated mice with cisplatin or aristolochic acid (AA) and stained for mouse IL-22 RNA and the PTC marker lotus lectin (LTL). Similar to the human samples, both cisplatin and AA induced expression of IL-22 RNA in mouse kidneys (Figure 2B). To confirm IL-22 gene expression in injured kidneys, we bred IL-22Cre mice with a floxed-stop-tdTomato reporter mice, to generate mice that express tdTomato under the control of the IL-22 promotor. Injuring IL-22Cre;tdTomato mice with cisplatin or AA injection induced tdTomato expression in kidney tubule cells (Supplemental Figure 1A).

**Figure 1.**
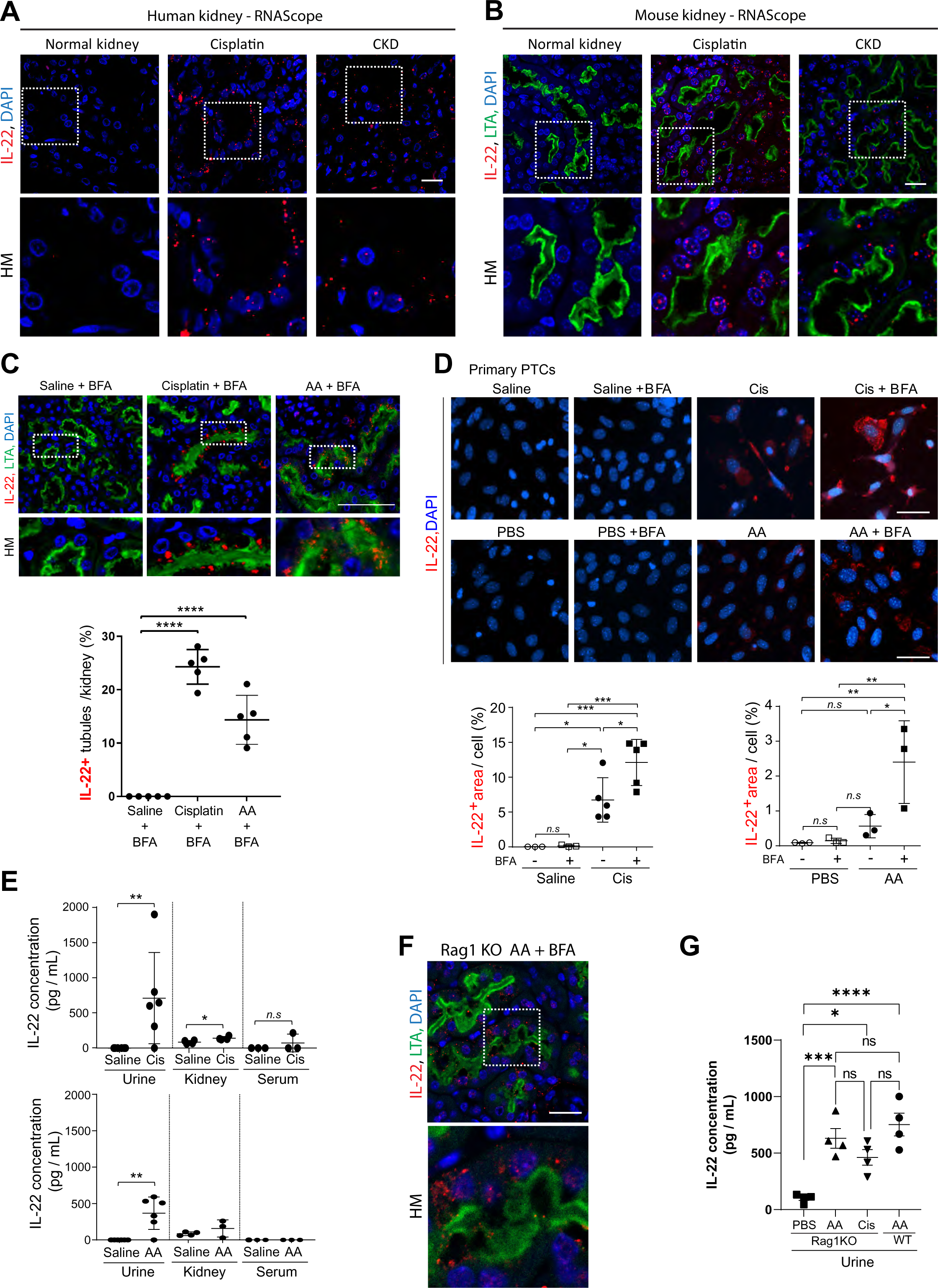
PTCs are a novel source of IL-22. (A) Representative images of IL-22 RNA staining using RNA scope in patient biopsies from patients diagnosed with cisplatin nephropathy or CKD. Scale bar: 20 µm. (B) Representative images of IL-22 RNA staining using RNA scope mouse tissue following cisplatin or AA induced AKI. Scale bar: 50 µm. (C) Representative immunofluorescence images from mouse kidneys following treatment with PBS + BFA, cisplatin + BFA, or AA + AA. Scale bar = 100 µm. (D) Representative immunofluorescence images from primary cultured PTCs treated with Saline, PBS, cisplatin, or AA + BFA. Scale bar: 50 µm. (E) IL-22 concentration measured by ELISA in mouse urine, kidney lysate, or serum at day 3 following saline, cisplatin, or AA injection. n = 3-6. (F) Representative immunofluorescence images of IL-22 in Rag1 KO mouse kidney following AA induced kidney injury and BFA injection. Scale bar: 20 µm. (G) IL-22 concentration measured by ELISA in Rag1 KO mice following PBS, cisplatin, or AA injection compared to wild-type treated with AA. Data are presented as the mean ±SD. Unpaired, two-tailed t-test (E) and one-way ANOVA and subsequent *Tukey’s post-hoc* test (C, D and G) were performed to identify statistical difference. *P<0.05, **P<0.01, ***P<0.001, and ****P<0.0001. AA, aristolochic acid. WT, wild-type.

**Figure 2.**
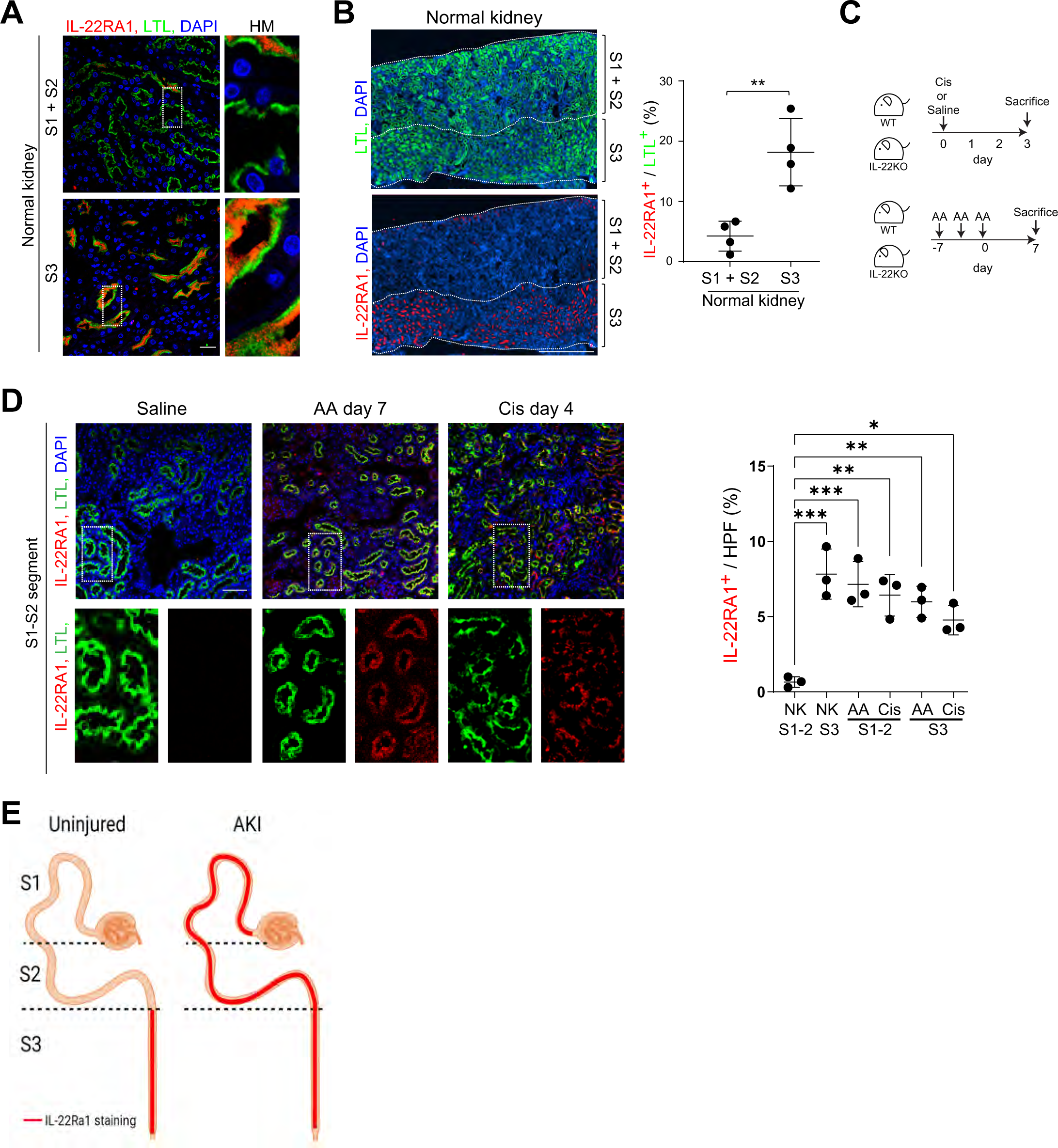
PTC IL-22RA1 expression is increased with injury. (A) Representative image of IL-22RA1 and LTL in S1-2 and S3 segments of normal kidney cortex. Scale bar: 20 μm. (B) Large scan image of IL-22RA1- and LTL-labeled normal kidney and quantification of IL-22RA1^+^ / LTL^+^ (%) in S1-2 and S3 segments of normal kidney cortex. n = 4, respectively. Scale bar: 500 μm. (C) Schematic diagram of cisplatin AKI and AA acute phase. (D) Representative images of IL-22RA1 expression in S1-S2 PTC segments following AA or cisplatin induced injury and quantification of IL-22RA1 positive area/HPF (%). N = 3. Scale bar: 50 μm. (E) Schematic diagram of the changes in IL-22RA1 expression following injury. Data are presented as the mean ±SD. Unpaired, two-tailed t-test was performed to identify statistical difference. **P<0.01. AA, aristolochic acid.

As IL-22 is a secreted protein, it is difficult to detect IL-22 protein within the source cells. To overcome this challenge, we injected mice with the secretion inhibitor brefeldin A (BFA) 6 hours before sacrifice and stained for IL-22 protein as shown previously^30^. Following cisplatin + BFA or AA + BFA treatment, IL-22 was detected in lotus lectin (LTL) positive PTCs (Figure 1C). To confirm PTC production of IL-22, we isolated PTCs from mice and treated with cisplatin or AA with/without BFA and stained for IL-22. Primary PTCs upregulated IL-22 in response to both cisplatin and AA, and the staining was enhanced with BFA (Figure 1D). The next question became, where is PTC IL-22 secreted? To answer this question, we performed ELISA analysis on serum, urine, and kidney lysates. Cisplatin or AA upregulated IL-22 predominantly in the urine (Figure 1E). To confirm the urinary IL-22 was from PTCs, we induced kidney injury in immune deficient Rag1 knockout (KO) mice, which lack mature T cells and B cells, using cisplatin or AA. Rag1 KO mouse kidneys stain positive for IL-22 in PTCs following AA + BFA injection (Figure 1F). Rag1 KO mice expressed a similar level of urinary IL-22 compared to wild-type mice (Figure 1G). Serum IL-22 was not increased in Rag1 KO mice (data not shown). These data suggest that PTCs are a novel source of IL-22 and contribute to urinary IL-22 levels.

### PTC IL-22RA1 expression is increased with injury

Given the high urinary expression of IL-22, we next sought to identify which kidney cell type presented the IL-22 receptor to the urine in uninjured mouse kidneys and kidneys following AKI. Immunofluorescent staining of IL-22RA1 revealed it was expressed on the brush boarder of PTCs, with staining most apparent in the S3 segment of the proximal tubule in uninjured kidneys (Figure 2A and 2B). Both cisplatin and AA induced injury upregulated IL-22RA1 in the S1 and S2 segments (Figure 2C, 2D, 2E). IL-22RA1 was also upregulated at the RNA level following injury (Supplemental Figure 1B). Co-staining for IL-22RA1 and the kidney injury marker KIM-1 revealed expression was lower in KIM-1+ PTCs (Supplemental Figure 1C). These data indicate that IL-22RA1 is constitutively expressed on the apical surface of S3 segment PTCs and is express in S1 + S2 segments with injury.

### IL-22 knockout mice are protected against AKI

To evaluate the role IL-22 in AKI, we treated wild-type and IL-22 global knockout mice (IL-22KO) with cisplatin and AA (Figure 3A) and followed mice over time. No IL-22 RNA was detected in IL-22KO mice with or without injury (Supplemental Figure 1D). Kaplan-Meier analysis showed cisplatin administration led to significantly less mortality in IL-22KO mice compared to WT mice (Figure 3B). Renal dysfunction, assessed by plasma creatinine, and weight loss were attenuated in IL-22KO in cisplatin and AA AKI (Figure 3C, 3D and Supplemental Figure 1E and 1F). Kidney injury molecule-1 (KIM-1), a marker of tubular injury, was upregulated in wild-type and IL-22KO mice after cisplatin or AA injury but the increase was significantly less in IL-22KO (Figure 3E and F). Similarly, the RNA levels of KIM-1, neutrophil gelatinase-associated lipocalin (NGAL), and IL-6 were significantly lower in IL-22KO mice following injury than wild-type (Supplemental Figure 1G, 1H). Cleaved caspase3 (C-casp3), a marker for apoptosis, was reduced in IL-22KO mice compared to wild-type following cisplatin or AA induced injury (Figure 3G-3H and Supplemental figure 1I). PTC differentiation marker Na/K-ATPase transporter was better preserved in injured IL-22KO mice compared to WT mice (Figure 3I-3J). PAS staining demonstrated kidney structure was better preserved in IL-22KO mice (Supplemental Figure 1J). Taken together, these data suggest deletion of IL-22 prevents AKI induced by cisplatin or AA.

**Figure 3.**
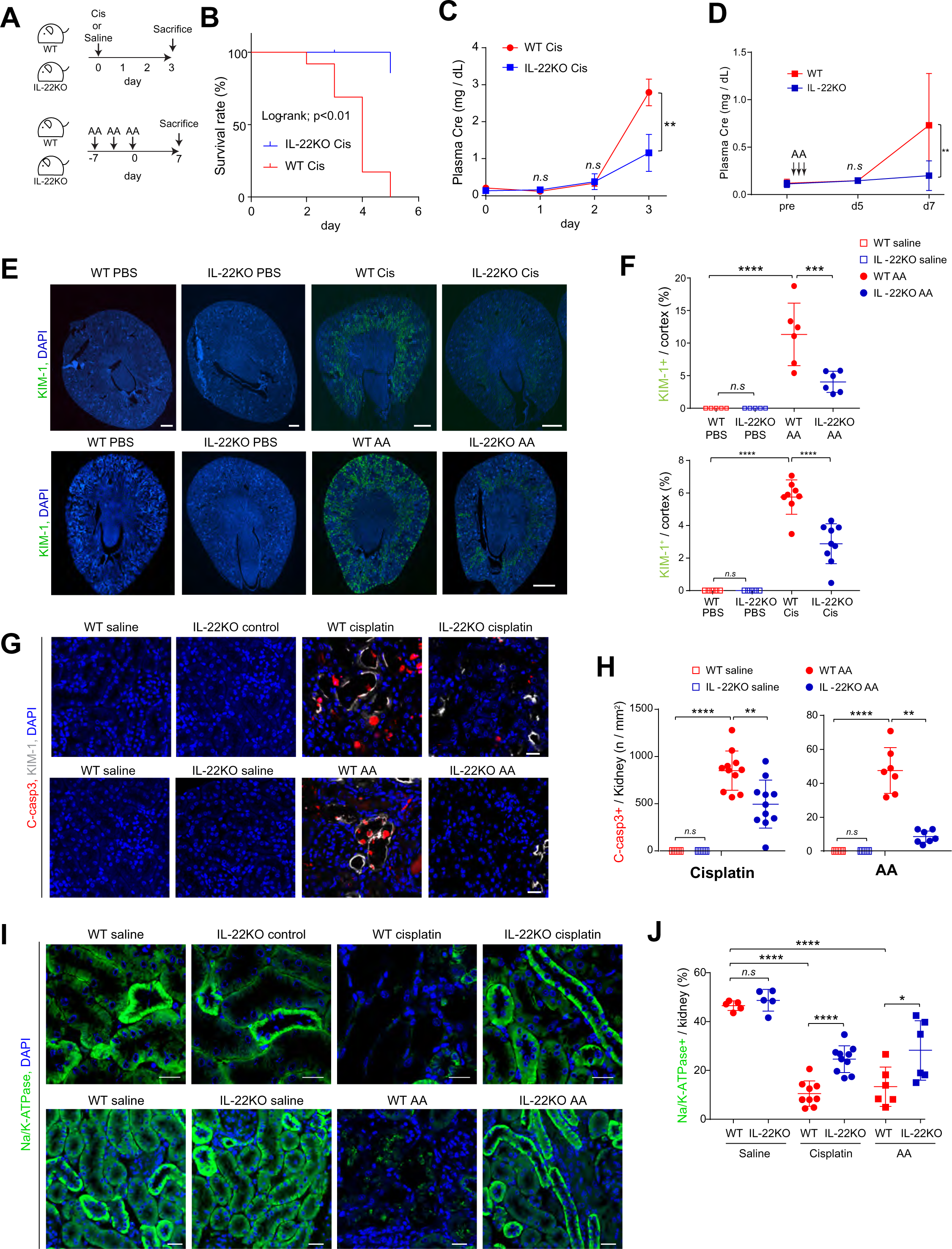
IL-22 knockout mice are protected against AKI. (A) Schematic diagram of cisplatin and AA induced AKI. (B) Survival rate in WT or IL-22KO mice after cisplatin injection. WT (n = 13) and IL-22KO (n = 10). (C) Plasma creatinine concentration over time after cisplatin or saline administration: WT (n = 12) and IL-22KO (n = 6). (D) Plasma creatinine on day 7 in AA. n = 5-12. (E) Representative images of KIM-1 staining in WT and IL-22 KO mice treated with PBS or AA. Scale bar: 500 µm (F) Quantification of the KIM-1 staining in F. KIM-1+ area/cortex area (%). n = 6. (G) Representative images of cleaved caspase 3 (C-casp3) in cisplatin AKI and AA acute phase. Scale bar: 20 μm. (H) Quantitative analysis for C-casp3+ cells / kidney (n / mm^2^). n = 5–11. (I) Representative images of NaKATPase staining in WT or IL-22KO kidneys following cisplatin or AA induced injury or vehicle control. Scale bar: 20 μm. (J) Quantitative analysis of NaKATPase+ area / kidney area (%). n = 5-9. Data are presented as the mean ±SD. Unpaired, two-tailed t-test (C and D) and one-way ANOVA and subsequent *Tukey’s post-hoc* test (F, H, and J) were performed to identify statistical difference. *P<0.05, **P<0.01, ***P<0.001, and ****P<0.0001. Cis, cisplatin; WT, wild-type; AA, aristolochic acid.

In addition to acute injury, AA is known to induce chronic kidney disease (CKD) and progressive fibrosis. We also examined the effect of IL-22 knockout on the chronic phase of AA induced injury. Chronic injury, as measured by serum creatinine, and weight loss up to 42 days post injury, were significantly reduced in IL-22KO compared to wild-type (Supplemental Figure 2A, 2B). Renal fibrosis, as measured by picrosirius red staining, was also significantly reduced in IL-22KO mice compared to WT mice (Supplementary Figure 2C). Consistent with a reduction in kidney injury, less immune cell infiltration was observed in IL-22KO mice following AA injury (Supplemental Figure 2D). These data suggest that the protective effect of IL-22 deletion in the acute phase is associated with reduce chronic injury.

### IL-22 promotes DDR activation

To investigate the mechanism by which IL-22 worsens kidney injury, we analyzed the phosphorylation and activation of DDR sensors H2AX and ATM in injured kidneys. The number of γH2AX^+^ cells, a specific marker of DNA damage, was increased in both WT and IL-22KO mice in cisplatin AKI and AA. However, the number of

γH2AX^+^ cells was significantly lower in AA-injured IL-22KO mice (Figure 4A). γH2AX^+^ cells were also reduced at 42 days post AA injection (Supplemental Figure 3A-3B). p-ATM^+^ cells were reduced in IL-22KO mice following cisplatin or AA exposure (Figure 4B). STAT3, one of the primary IL-22/IL-22RA1 signaling molecules, was upregulated in both injured wild-type and IL-22KO, but to a lesser extent in the IL-22KO group (Figure 4C). Phosphorylated p53 (p-p53) and total p53 were increased in WT kidney of cisplatin AKI, which were reduced in IL-22KO mice in parallel with MDM2 levels (Figure 4C-4D and Supplemental Figure 3C-3D). These data suggest that IL-22 upregulates DDR activation *in vivo*.

**Figure 4.**
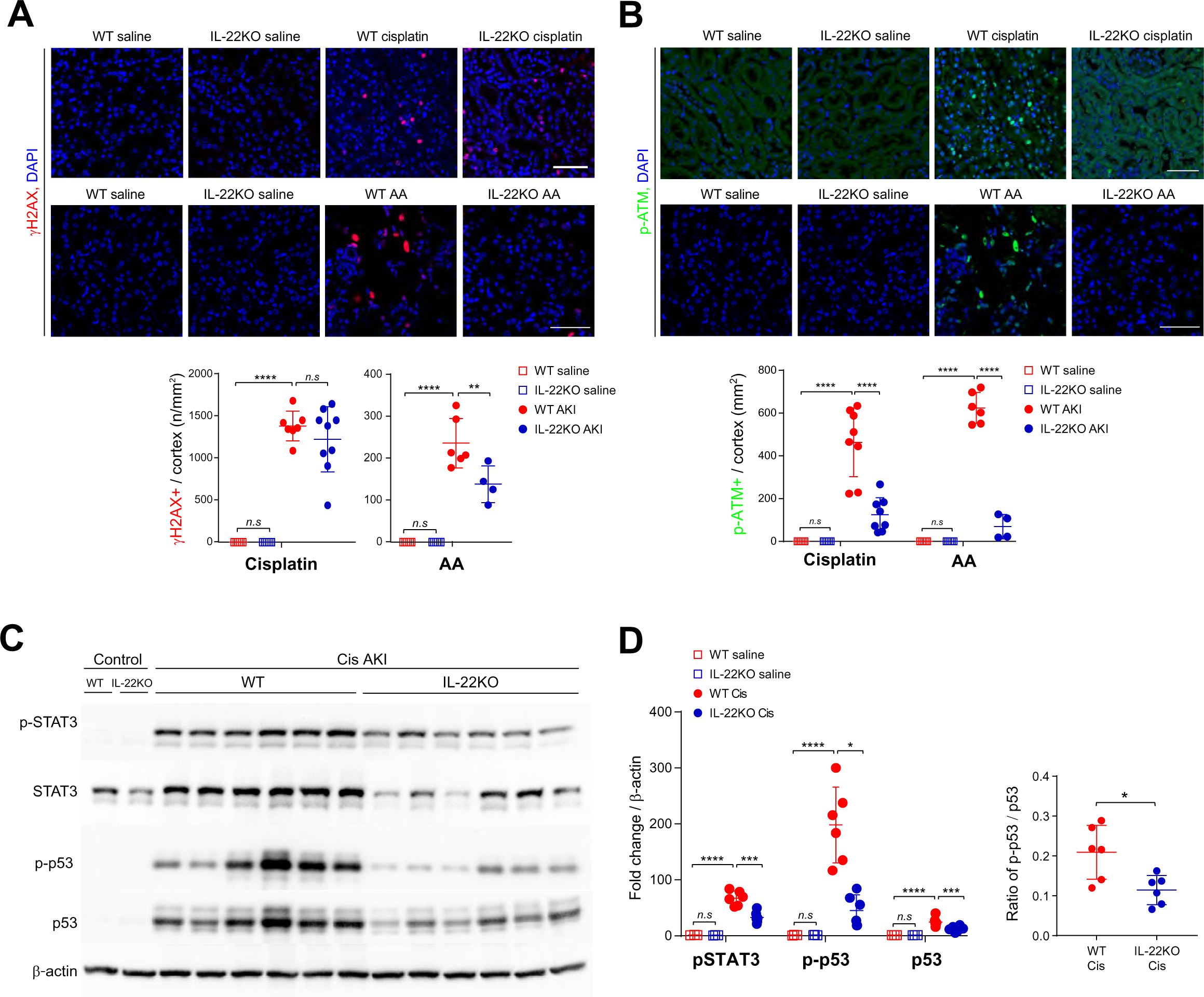
IL-22 promotes DDR activation. (A) Representative images of γH2AX-labeled kidney sections from WT and IL-22KO mice in cisplatin or AA treated mice (upper panel). The corresponding quantification of γH2AX^+^ cells (n / mm^2^) (lower panel). Scale bar: 50μm. n = 5–9. (B) Representative images of p-ATM-labeled kidney sections from WT and IL-22KO mice in cisplatin or AA treated mice (upper panel). The corresponding quantification of p-ATM^+^ cells (n / mm^2^) (lower panel). Scale bar: 50μm. n = 5–8. (C) Western blot analysis of pSTAT3, STAT3, p-p53, p53, and β-actin in kidneys of WT and IL-22KO mice on day 3 after administration of saline or cisplatin. (D) Quantitative analysis of pSTAT3/ β-actin, p-p53/ β-actin, p53/ β-actin, and p-p53/ p53. Data are presented as the mean ±SD. Unpaired, two-tailed t-test (D) and one-way ANOVA and subsequent *Tukey’s post-hoc* test (A, B, and D) were performed to identify statistical difference. *P<0.05, **P<0.01, ***P<0.001, and ****P<0.0001. Cis, cisplatin; WT, wild-type; AA, aristolochic acid; ATM, Ataxia-telangiectasia mutated; ATR, Ataxia-telangiectasia and rad3-related; PUMA, p53 upregulated modulator of apoptosis.

### Knockout of IL-22RA1 in kidney nephron protects against AKI

While our data suggests that IL-22 worsens AKI by acting on intrinsic kidney epithelial cells, it remains possible that it functions in an extra-renal manner. To confirm that IL-22 promotes kidney injury by directly acting on PTCs PTCs *in vivo*, we deleted IL-22RA1 specifically in kidney epithelial cells by crossing *IL-22RA1flox/flox* with *Six2Cre* (IL-22RA1^ΔTub^). *IL-22RA1*^ΔTub^ mice developed normally and had no observed kidney phenotype prior to injury. We confirmed that IL-RA1 deletion abolished IL-22 signaling in PTCs by treating primary IL-22RA1^ΔTub^ PTCs with PBS or IL-22 +/- cisplatin or AA. IL-22 incubation did not promote STAT3 phosphorylation, a primary target of IL-22RA1 (Supplemental Figure 4A). IL-22RA1^ΔTub^ were injected with cisplatin to induce AKI (Figure 5A). Immunostaining revealed that IL-22RA1 expression was significantly reduced in kidney of *IL-22RA1*^ΔTub^ mice (Figure 5B-5C). Following cisplatin injection, renal functional decline was attenuated in the IL-22RA1^ΔTub^ mice (Figure 5D). PAS staining demonstrated kidney structure was better preserved in IL-22KO mice (Supplemental Figure 4B). Immunostaining revealed c-casp 3 levels were increased in IL-22RA1fl/fl mice but not IL-22RA1^ΔTub^ (Figure 5E-5F). The number of gH2AX+ and p-ATM+ cells were reduced in IL-22RA1^ΔTub^ mice compared to IL-22RA1fl/fl following cisplatin exposure (Figure 5G-5J). Similar to IL-22KO mice, pSTAT3 and p-53 levels were upregulated in control mice following cisplatin injection, but reduced in IL-22RA1^ΔTub^ (Figure 5K, Supplemental Figure 4C). Taken together, these data indicate that IL-22 signaling via IL-22RA1 in kidney epithelial cells enhanced DDR activation and kidney injury in AKI.

**Figure 5.**
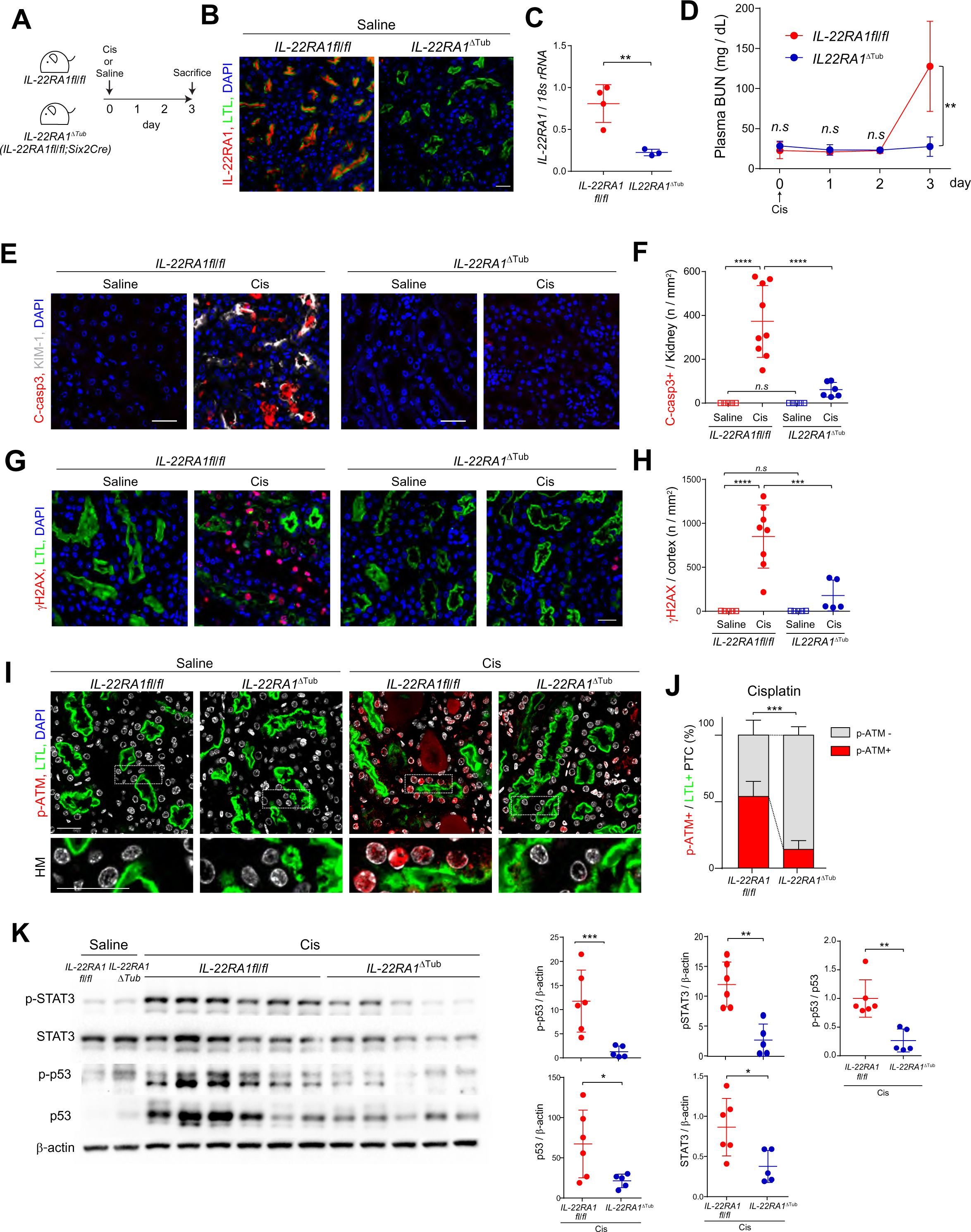
Knockout of IL-22RA1 in kidney nephron protects against AKI. (A) Schematic diagram of cisplatin AKI with IL-22RA1^ΔTub^ and IL-22RA1fl/fl. (B) Representative image of IL-22RA1 and LTL in S3 segments of IL-22RA1^Δ^ ^Tub^ and IL-22RA1fl/fl. Scale bar: 50 µm. (C) Real time PCR for *IL-22RA1* in kidneys of IL-22RA1^Δ^ ^Tub^ and IL-22RA1fl/fl. n = 3-4. (D) Plasma BUN level over time after cisplatin or saline injection. (E) Representative images of C-casp3 in kidneys of IL-22RA1^Δ^ ^Tub^ and IL-22RA1fl/fl in cisplatin AKI. Scale bar: 20 μm. (F) Quantitative analysis for C-casp3+ cells / kidney (n / mm^2^). n = 5–6. (G) Representative images of γH2AX staining in IL-22RA1^Δ^ ^Tub^ and IL-22RA1fl/fl in cisplatin AKI. Scale bar: 20 μm. (H) Quantitative analysis of γH2AX+ cells / kidney (n / mm^2^). n = 5–8. (I) Representative images of p-ATM and LTL-stained IL-22RA1fl/fl or IL-22RA1^ΔTub^ kidneys treated with saline or cisplatin. Scale bars: 20 µm. (J) The corresponding quantification of p-ATM+ nuclei in PTC. n = 5, respectively. (K) Western blots for pSTAT3, STAT-3, p-p53, p53 and β-actin in kidneys from in kidneys from *IL-22RA1*^Δ^ ^Tub^ and *IL-22RA1fl/fl* following cisplatin injury. Data are presented as the mean ±SD. Unpaired, two-tailed t-test (C, D, J, and K) and one-way ANOVA and subsequent *Tukey’s post-hoc* test (F and H) were performed to identify statistical difference. *P<0.05, **P<0.01, ***P<0.001, and ****P<0.0001. ATM, Ataxia-telangiectasia mutated; ATR, Ataxia-telangiectasia and rad3-related; PUMA, p53 upregulated modulator of apoptosis.

### IL-22 enhances DDR activation in vitro

To confirm whether IL-22 induces DDR activation in PTCs, we treated immortalized mouse renal proximal tubule epithelial cells (RPTEC) (which express a high level of IL-22 receptor, IL-22RA1 (Supplemental Figure 4D)) with different concentrations of recombinant (rIL-22). Treatment with rIL-22 for 15 mins upregulated p-STAT3, p-p53, and p-MDM2(Figure 6A, Supplemental Figure 4E). Combining rIL-22 with cisplatin upregulated p-STAT3 more than either agent alone (Figure 6B). At longer time points, c-casp3 was induced by 24 hours incubation with rIL-22 alone (Figure 6C). Co-incubation of rIL-22 with a selective STAT3 inhibitor, S3I-201, prevented the phosphorylation of p53 and activation of caspase 3 (Figure 6C-6E). These data suggests that IL-22 regulation of STAT3 is necessary for IL-22 induced p53 activation and apoptosis.

**Figure 6.**
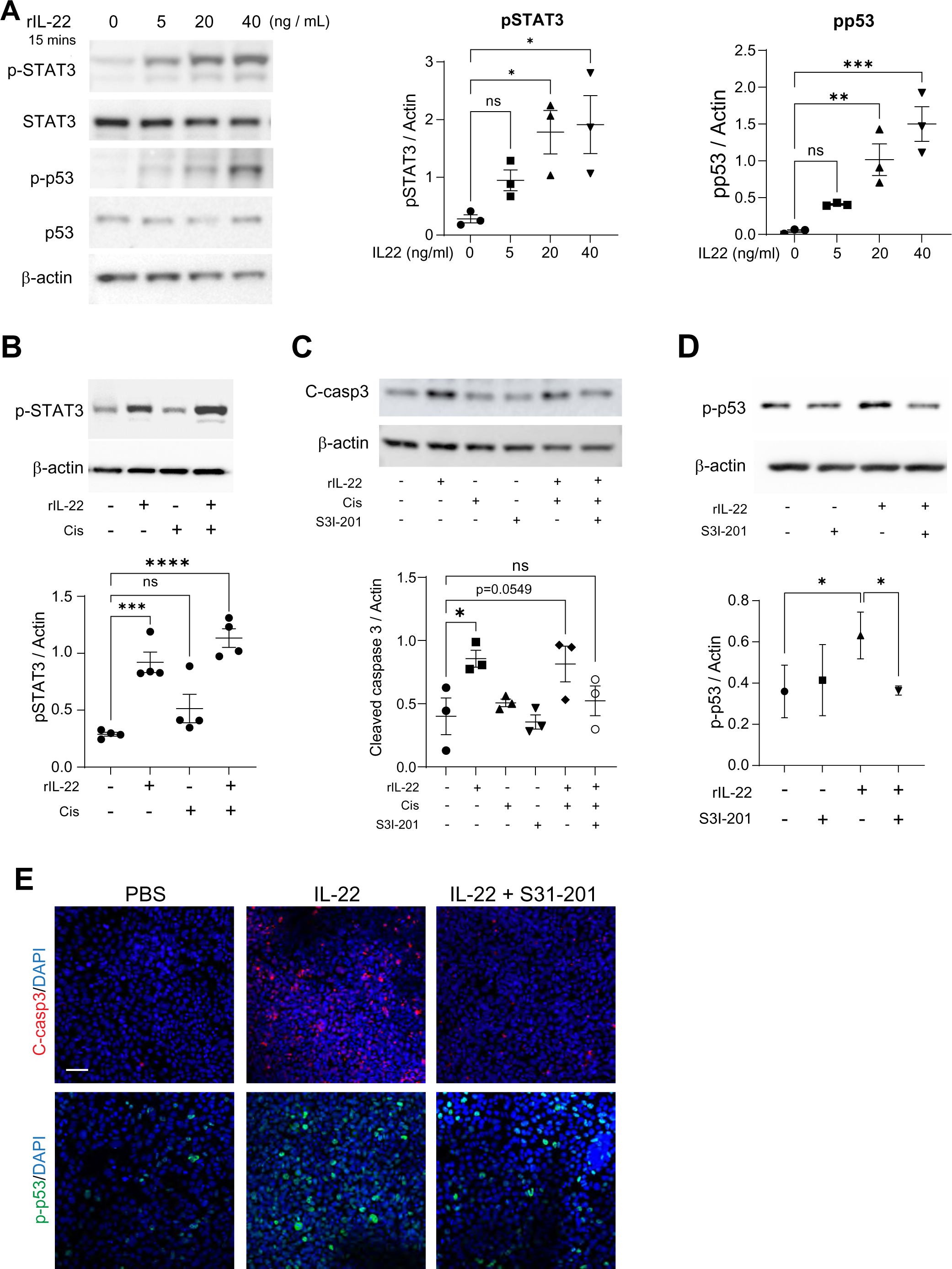
IL-22 enhances DDR activation in vitro. (A) Western blot and quantitative analysis of pSTAT3, STAT3, p-p53, p53, and β-actin in RPTEC incubated with escalating dose of rIL-22 for an hour. (B) Western blot and quantitative analysis of pSTAT3 in RPTEC treated with cisplatin with or without rIL-22. (C) Western blot and quantitative analysis of C-casp3 in RPTEC incubated with rIL-22 (10 ng / mL) for 24 hours in the presence or absence of S3I-201. (D) Western blot and quantitative analysis of p-p53 and β-actin in RPTEC incubated with rIL-22 with or without S3I-201 for an hour. (E) Representative immunofluorescence images of RPTCs treated with PBS, IL-22 or IL-22+S31-201 and stained for cleaved caspase 3 or p-p53. Scale bar = 50 µm. C-Casp3, cleaved caspase 3; Cis, cisplatin; WT, wild-type; AA, aristolochic acid.

Fusion proteins consisting of IL-22 fused to Fc region of IgG or whole antibodies have been developed as therapeutic agents, with IL-22-Fc entering into clinical trials^31^. Previous studies have found that IL-22-antibody fusion proteins protect against kidney injury, through regulation of the NOD-, LRR- and pyrin domain-containing protein 3 (Nlrp3) inflammasome or mitochondrial protective functions^25, 32–36^. It should be noted, however, that adding an Fc tag alters IL-22 function, and IL-22-Fc can act as an antagonist^37^. To evaluate whether untagged rIL-22 or IL-22-Fc regulated inflammasome activation in our model, we treated IL-22KO primary PTCs with AA +/-rIL-22 or IL-22-Fc and analyzed the expression of Nlrp3 and its adaptor protein apoptosis-associated speck-like protein containing a caspase-recruitment domain (ASC). The combination of rIL-22 + AA did not alter the expression of Nlrp3 RNA but did increase the expression of ASC (Supplemental Figure 5A). Conversely, the combination of IL-22-Fc + AA reduced both Nlrp3 and ASC RNA compared to AA alone (Supplemental Figure 5A). To test inflammasome activation, we analyzed the cleavage of caspase 1, an important step in inflammasome activation^38^. The combination of rIL-22 and AA increased caspase 1 and cleaved caspase 1 levels, while IL-22-Fc reduced caspase 1 cleavage (Supplemental Figure 5B). These results suggest IL-22-Fc suppresses Nlrp3 inflammasome, similar to previously reports, while untagged IL-22 does not. Thus, while IL-22-Fc is a promising therapeutic, it may not be a representative tool to study IL-22 biological functions.

### IL-22 activates the DDR through STAT3-ATM signaling

As our results indicate PTCs can produce IL-22, we evaluated the effects of rIL-22 supplementation on in IL-22KO primary PTCs. To determine if IL-22 amplifies PTC DDR activation, primary PTCs were treated with cisplatin and/or rIL-22 and analyzed for upregulation of DDR components, p53, ATM, and PUMA (a pro-apoptotic p53 target), and cell death. Cisplatin alone induced a marginal increase in p53, while the combination of cisplatin + rIL-22 upregulated p53 in both wild-type and IL-22KO PTCs (Figure 7A). Similarly, the combination of AA + rIL-22 upregulated c-caspase 3 (Figure 7B). rIL-22 also upregulated γH2AX levels in cisplatin or AA treated PTCs (Figure 7C). Interestingly, treatment with IL-22-Fc + cisplatin or AA did not increase γH2AX above cisplatin or AA alone (Supplemental Figure 5C). AA treatment upregulated *ATM* and *PUMA* at the mRNA level, which was further enhanced by rIL-22 (Figure 7D-7E).

**Figure 7.**
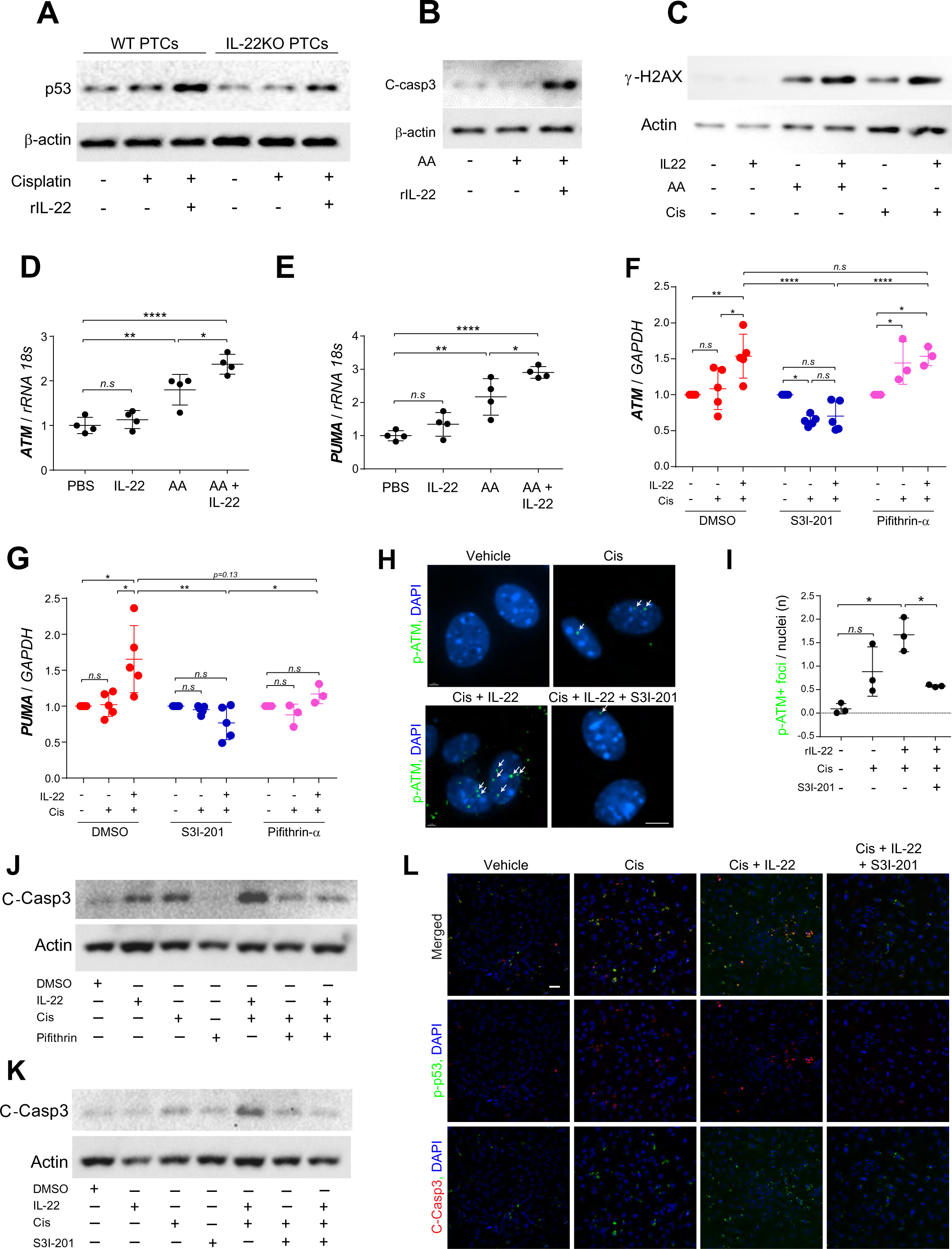
IL-22 activates the DDR in primary PTCs. (A) Western blot analysis of p53 in primary PTCs from WT and IL-22KO mice treated with cisplatin in the presence or absence of rIL-22. (B) Western blot analysis for C-casp3 in IL-22KO primary cells treated with AA in the presence or absence of rIL-22. (C) Western blot analysis of yH2AX levels in IL-22KO primary PTCs following cisplatin or AA treatment +/-rIL-22. (D-E) Real-time PCR analysis of ATM and PUMA in IL-22KO primary PTCs treated with AA in the presence or absence of rIL-22 for 7 days. (F-G) Real-time PCR analysis of ATM and PUMA in IL-22KO primary PTCs treated with cisplatin or cisplatin plus rIL-22 in the presence of S3I-201, pifithrin-α, or DMSO (vehicle). (H) Representative images of p-ATM foci in IL-22KO primary PTCs treated with cisplatin or cisplatin plus rIL-22 in the presence or absence of S3I-201. Scale bar: 3μm. (I) The corresponding quantification of number of p-ATM+ foci in nuclei (n / nuclei). n = 3, respectively. (J) Western blot analysis of C-casp3 in primary PTCs treated with cisplatin, IL-22 and/or pifithrin-α. (K) Western blot analysis of C-casp3 in primary PTCs treated with cisplatin, rIL-22 and/or S31-201. (L) Immunofluorescence staining of p-p53 and C-casp3 in primary PTCs treated with cisplatin, rIL-22 and/or S31-201. Scale bar: 20 µm. Data are presented as the mean ± SD. One-way ANOVA and subsequent *Tukey’s post-hoc* test was conducted to identify statistical difference. *P<0.05, **P<0.01, ***P<0.001, and ****P<0.0001. Cis, cisplatin; WT, wild-type; AA, aristolochic acid; ATM, Ataxia-telangiectasia mutated; ATR, Ataxia-telangiectasia and rad3-related; PUMA, p53 upregulated modulator of apoptosis.

To investigate if the DDR activation is mediated by IL-22-induced STAT3 activation, IL-22KO primary PTCs were treated with cisplatin and/or rIL-22 in the presence of STAT3 inhibitor S3I-201 or p53 inhibitor pifithrin-α. rIL-22-induced ATM upregulation was inhibited by S3I-201, but not pifithrin-α, meanwhile the upregulation of PUMA (a downstream target of p53) was suppressed by both S3I-201 and pifithrin-α (Figure 7F-7G). S3I-201 also reduced the number of nuclear p-ATM foci in IL-22KO primary PTC treated with cisplatin in the presence of rIL-22 (Figure 7H-7I). Treating PTCs with either rIL-22 or cisplatin induced an upregulation of c-caspase 3, while the combination of the two further increased c-caspase 3. Inhibition of p53 with pifithrin prevents cisplatin and rIL-22 induced caspase 3 activation (Figure 7J-7L). Similarly, inhibition of STAT3 with S3I-201 prevents caspase 3 activation and reduces p-p53 levels (Figure 7K-7L). Taken together, these data indicate that IL-22 enhances DDR activation and PTC injury from cisplatin or AA exposure, and inhibition of either STAT3 or p53 can inhibit the injury.

## Discussion

The goal of this study was to identify the kidney specific role of IL-22 in AKI and examine its role in regulation of the DDR in kidney epithelial cells. We demonstrate new findings as follows: (1) PTCs are a novel source, and to our knowledge the only epithelial source, of IL-22; (2) IL-22 global knockout prevents AKI induced by cisplatin or AA; (3) IL-22 induced injury is kidney intrinsic, as kidney specific deletion of the IL-22 receptor also ameliorates AKI; (4) IL-22 amplifies DDR signaling, including ATM, p53, and PUMA, promoting apoptosis; (5) Inhibition of STAT3 or p53 downstream of IL-22 mimics the anti-apoptotic phenotype of IL-22 and IL-22RA1 knockouts. We demonstrate that IL-22/IL-22RA1 binding triggers rapid phosphorylation of STAT3. ATM is then upregulated in a STAT3-dependent manner (Figure 8). These findings are supported by Gronke et al. that showed STAT3 binds to promoter region of ATM^18^. ATM can then directly phosphorylate p53 and MDM2 to initiate the downstream pro-apoptotic factors including PUMA^39^. While IL-22 induced DDR is not enough to cause kidney injury under basal condition, in the setting of drugs that induce significant DNA damage, e.g. cisplatin or AA, IL-22 can further activate the DDR and promote apoptosis (Figure 8). Thus, IL-22 amplifies the PTC DDR signaling and shifting the cell fate to apoptosis.

**Figure 8.**
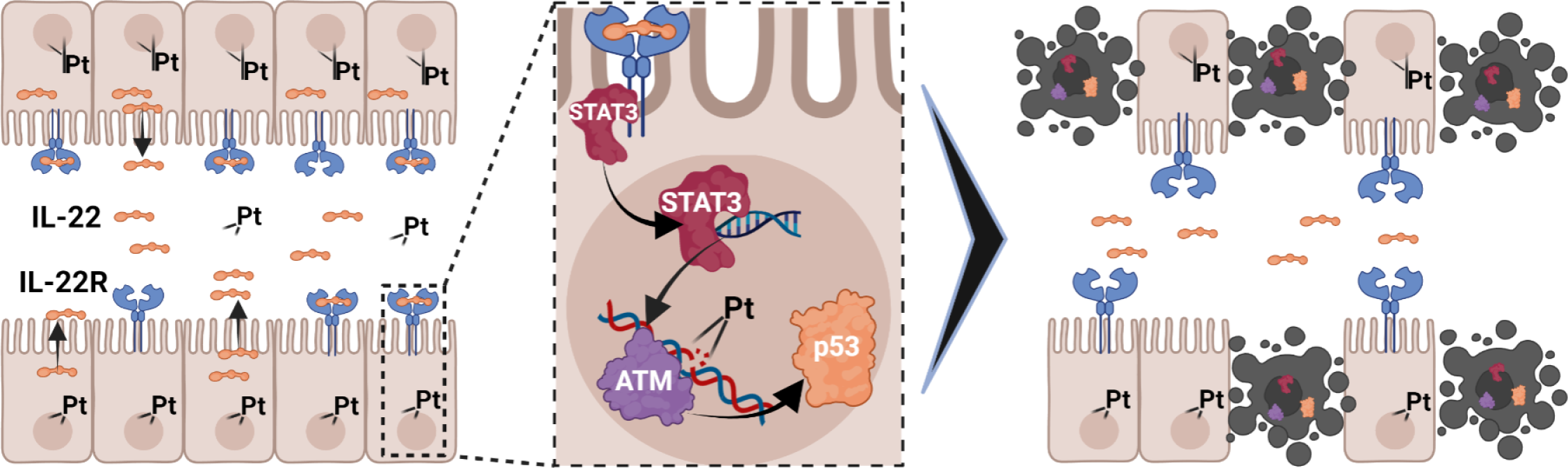
Proposed mechanism of IL-22 induced DDR activation. Cisplatin or AA-induced injury induces IL-22 secretion from PTCs, activating IL-22RA1. IL-22/IL-22RA1 activates STAT3 to promote transcription of ATM. ATM then phosphorylates p53. In combination with cisplatin induced injury, further upregulation of DDR by IL-22 promotes cell death.

Overactivation of STAT3 and the DDR have been linked to AKI in animal models and patient samples^2, 3, 40–45^. Despite this fact, the importance of these pathways in maintaining homeostasis has limited the development of therapeutics targeting the pathways directly. Here we demonstrate a novel mechanism to prevent the overactivation of these pathways. The current paradigm suggests the outcome of DDR activation is directly proportional to the level of DNA damage, with mild DNA damage leading to DDR mediated repair and cell survival and severe damage promoting DDR directed cell death^46–48^. Our findings challenge this paradigm by demonstrating extracellular signaling via IL-22 shifts the balance towards cell death. While deletion of IL-22 or IL-22RA1 does not prevent DNA repair by the DDR, it does prevent overactivation of STAT3 and DDR beyond the drug induced injury alone. This pushes already injured PTCs towards cell death and worsens nephrotoxic AKI. Thus, this study provides the first mechanistic insight into how IL-22 dictates life/death decisions in injured PTCs and demonstrates a novel approach to target STAT3 and p53 overactivation in AKI without interfering with their basal function.

Another novel finding of the current study is that PTCs are a previously unrecognized source of IL-22. To our knowledge, no other epithelial cells have been reported to produce IL-22. While this is a surprising result, it is consistent with the highly immunogenic transformation injured PTCs undergo. PTCs are known to become semiprofessional antigen presenting cells which upregulate immune receptors, including KIM-1 (also known as T-cell, Ig and mucin containing 1 (TIM-1)) and MHCII, in response to injury^49, 50^. KIM-1 expression in turn transforms PTCs into phagocytes^51^. PTCs are also known to secrete a number of cytokines, including TGFβ, TNFα, IL-6, IL-10 and etc^52^. The primary function of IL-22 is strengthening epithelial barrier integrity and other anti-bacterial responses, and IL-22 signaling is an important component of the antibacterial response in bladder epithelial cells^53^. This suggests PTC secretion of urinary IL-22 is likely a physiological response to stress, such as urinary tract infections, but this process becomes active in AKI as well. Given the fact that PTCs are the only cell type in the kidney to express IL-22RA1in the kidney, however, in the setting of kidney injury, there is autocrine/paracrine IL-22 signaling within the PT. While this paracrine signaling may be helpful in the setting of infection, our evidence suggests it is detrimental in nephrotoxin induced AKI.

Drug induced nephropathy (DIN) is a major health concern, accounting for over 60% of AKI hospitalizations in the elderly population ^3, 54–56^. As the population ages, DIN is expected to become more common. Herbal remedies and nutritional supplements which contain nephrotoxic compounds, such as AA, can further complicate DIN^57, 58^. While some nephrotoxic drugs can be substituted for less toxic alternatives, many nephrotoxic drugs, such as cisplatin used as first-line therapy for many tumors, do not have non-nephrotoxic alternatives. Here we demonstrate that drugs which induce kidney injury through DNA damage are better tolerated and induce less kidney injury without IL-22 signaling. These data suggest that IL-22 inhibition is a potential therapeutic strategy to prevent DIN from these drugs.

IL-22 signaling is known to result in pro-reparative or pathological response, depending on the context. IL-22’s ability to strengthening epithelial barrier integrity is thought to provide protection against acute injuries, such as ischemic, acetaminophen, and alcohol injuries to the liver through accelerated wound healing^15–17^. On the other hand, IL-22 promotes injury in irritable bowel disease, Crohn’s disease, dermatitis, and lupus nephritis^20–22^. Recent evidence suggests the use of different IL-22 dosages and reagents can contribute to contradictory results. Endogenous IL-22 serum concentration rarely surpasses 200 pg/ml; however, studies often use animal models with IL-22 serum concentrations between 600-7,000 pg/ml^59, 60^. Studies in obesity models have demonstrated treatment of mice with very high doses of IL-22 or IL-22-Fc reduced liver and body fat, while endogenous levels did not affect fat deposition^61, 62^. IL-22 fusion proteins can also cause different results. For instance, while IL-22-Fc has been shown promise as a therapeutic, it activates different cellular signaling compared to endogenous or untagged IL-22 ^37^. N-terminally tagged Fc-IL-22 has been shown to amplify IL-22 signaling, while C-terminally tagged IL-22-Fc acts as an antagonist^63^. In the kidney, recent studies have shown that IL-22-antibody or IL-22-Fc fusion constructs protect against kidney injury, while untagged IL-22 does not^22, 25, 27, 32–36^. Similarly, we find IL-22-Fc inhibits inflammasome activation; however, untagged rIL-22 does not suppress inflammasome activation. Another limitation of the supplementation approach is that it targets multiple organs expressing IL-22RA1. To circumvent the issues listed above, we studied IL-22 using kidney tubule specific deletion of IL-22RA1 and global genetic IL-22 deletion. In both models, we found IL-22 signaling in PTCs promotes injury and death in AKI through overactivation of the DDR. As IL-22 fusion constructs show great therapeutic potential and have entered clinical trials, it is important to study these promising drug candidates; however, the fact they have been modified to optimize protective effects should also be considered. Therefore, models utilizing super-physiological IL-22 concentrations or IL-22-Fc may not be the ideal tools to study native IL-22 function and pathobiology.

## Conclusion

Here we have described a novel pro-injury and pro-cell death role of IL-22 in injured kidneys. PTCs represent a novel source of IL-22. IL-22/IL-22RA1 binding on PTCs amplifies the DDR response and, in combination with DNA damaging nephrotoxins, tips the life/death balance towards cell death. Global deletion of IL-22 or kidney specific deletion of IL-22RA1 reduces activation of DDR components ATM, p53, and pro-apoptotic PUMA. Therapeutics targeting IL-22 given alongside nephrotoxic drugs could improve the tolerance of these drugs and prevent kidney injury.

## Acknowledgements

We would like to thank Dr. Ray Harris, Dr. Roy Zent, and Dr. Ambra Pozzi for their help in manuscript preparation. We would like to thank Sushrut S. Waikar and Samir M Parikh for help obtaining biopsy samples. We would also like to acknowledge the Vanderbilt Cell Imaging Shared Resource and Nikon Center of Excellence (supported by NIH grants DK020593, CA68485, DK20593, DK58404, DK59637 and EY08126) for help with imaging and analysis.

## Author contributions

K.T.; writing–original draft preparation. B.E., E.S., N.P., and C.B.; writing–review and edit. K.T., B.E., and S.S. performed all experiments and analyzed the data. All authors have read and agreed to the final version of manuscript.

