## Supplementary Materials for "IL-22 promotes acute kidney injury through activation of the DNA damage response and cell death in proximal tubule cells"

Kensei Taguchi et al.

Supplemental Material Table of Contents:

Legends of Supplemental Figures

Supplemental Figure 1

Supplemental Figure 2

Supplemental Figure 3

Supplemental Figure 4

Supplemental Figure 5

Supplemental Table 1

Supplemental Table 2

Supplemental Methods

Supplemental References

**Supplementary Figure 1. IL-22 knockout protects kidney against AKI and subsequent CKD**

(A) IL-22Cre:tdTomato mice were injected with cisplatin or AA. Kidneys were sectioned and stained for tdTomato and NaKATPase. Scale bar = 20 μm. (B) Real-time PCR analysis of *IL-22RA1* in WT kidneys on day 7 and day 42 following AA exposure. Scale bar = 20 μm. (C) Representative images of IL-22RA1 and KIM-1 expression in PTC segments following AA or cisplatin induced injury. (D) Real-time PCR analysis of IL-22 mRNA in uninjured and injured kidneys from wild-type and IL-22KO mice. (E) Body weight (compared to day 0) on day 3 after injection of saline and cisplatin. n = 12-13. (F) Body weight (compared to day 0) after administration with AA. n = 5-12. (G) Real-time PCR analysis of KIM-1 and NGAL mRNA in uninjured and injured kidneys from wild-type and IL-22KO mice. (H) Real-time PCR analysis of IL-6 in uninjured and injured kidneys from wild-type an IL-22KO mice. (I) Western blot analysis of C-casp3 of kidneys of WT and IL-22KO on day 3 after saline or cisplatin administration. (J) Representative images of PAS stained wild-type and IL-22KO kidneys following PBS, cisplatin or AA injection. Scale bar: 100 μm. Data are presented as the mean ± SD. Unpaired, two-tailed t-test (E and F) and One-way ANOVA and subsequent *Tukey's post-hoc* test (B, D, G, and H) were performed to identify statistical difference. *P<0.05, **P<0.01, ***P<0.001, and ****P<0.0001. C-MDM2, cleaved Mouse double minute 2 homolog.

**Supplementary Figure 2. AA does not induce chronic kidney injury in IL-22KO mice.**

(A) Schematic diagram of AA chronic phase injury. (B) Plasma creatinine and body weight (compared to day 0) taken weekly until week 6 after AA administration. n = 4. (C) Representative images of picrosirius red staining in WT mice and IL-22KO mice on day42 and the corresponding quantitation of collagen+ / kidney (%). Scale bar: 500 μm. n = 4. (D) Representative immunofluorescence images of AA treated kidneys stained for F4/80 and CD3. Scale bar: 1 mm. n = 5. Data are presented as the mean ± SD. Unpaired, two-tailed t-test (B, C, and D) was performed to identify statistical difference. *P<0.05, **P<0.01, ***P<0.001, and ****P<0.0001. C-MDM2, cleaved Mouse double minute 2 homolog.

**Supplemental Figure 3. IL-22 amplifies DDR in AKI and CKD**

(A) Representative images of γH2AX- and p-ATM-labeled kidney from WT and IL-22KO mice on day 7 (acute) and day42 (chronic) of AA. Scale bar: 20 μm. (B) The corresponding quantification for number of γH2AX$^+$ cells on day 7 and day 42 in WT and IL-22KO. n = 5-9. (C) Western blots for C-MDM2 of WT and IL-22KO on day3 after saline or cisplatin administration. (D) The corresponding quantification of C-MDM2 / β-actin. Data are presented as the mean ± SD. Unpaired, two-tailed t-test (B) and One-way ANOVA and subsequent *Tukey's post-hoc* test (D) were performed to identify statistical difference. *P<0.05, **P<0.01, ***P<0.001, and ****P<0.0001. C-MDM2, cleaved Mouse double minute 2 homolog.

**Supplemental Figure 4. IL-22RA1 KO prevents DDR activation.**

(A) Western blot analysis of p-STAT3 activation in IL-22RA1 primary PTCs treated with rIL-22. (B) Representative images of PAS stained wild-type and IL-22RA1 $^{\Delta Tub}$ kidneys following PBS or cisplatin. Scale bar: 100 μm. (C) Western blot analysis of p-p53 and p53 in wild-type or IL-22RA1 $^{\Delta Tub}$ kidneys following AA administration. (D) Western blot analysis of IL-22RA1 expression in RPTEC cell line. (E) Western blot analysis of p-MDM2 in RPTECs treated with increasing doses of IL-22. Unpaired, two-tailed t-test (C) was performed to identify statistical difference. *P<0.05, **P<0.01, ***P<0.001, and ****P<0.0001

**Supplemental Figure 5. rIL-22 promotes inflammasome activation, while IL-22-Fc does not.**

(A) Real-time PCR analysis of NLRP3 and ASC levels in IL-22KO primary PTCs treated with cisplatin, AA and rIL-22 or IL-22-Fc. (B) Representative images of caspase 1 or cleaved caspase 1 staining in IL-22KO primary PTCs treated with AA + rIL-22 or IL-22-Fc. Scale bar: 50 μm. (C) Western blot analysis of γH2AX expression in IL-22KO primary PTCs treated with cisplatin, AA Data are presented as the mean ± SD. ANOVA and subsequent *Tukey's post-hoc* test (A) was performed to identify statistical difference. *P<0.05, **P<0.01, ***P<0.001, and ****P<0.0001

### Supplementary Figure 1

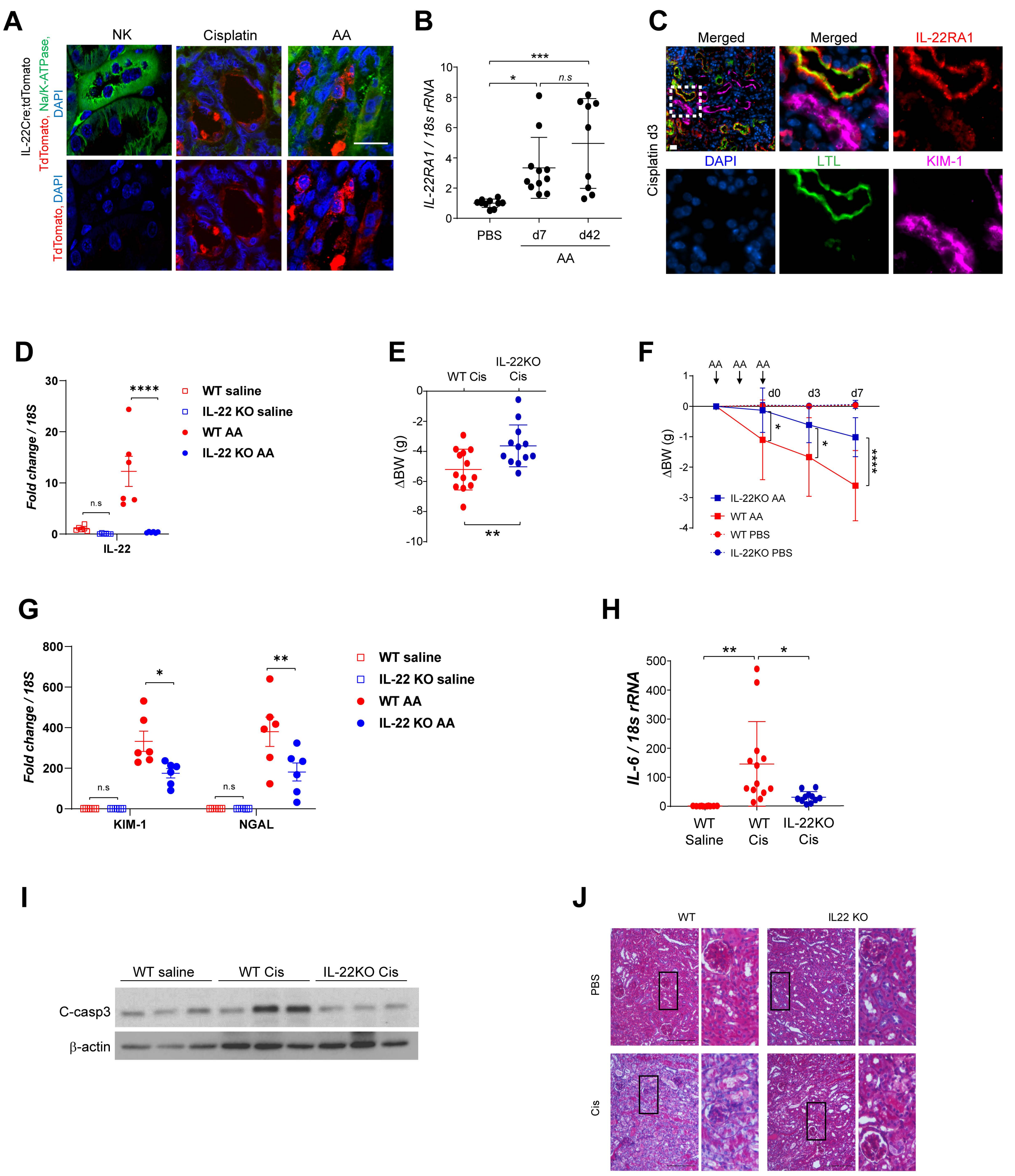

### Supplementary Figure 2

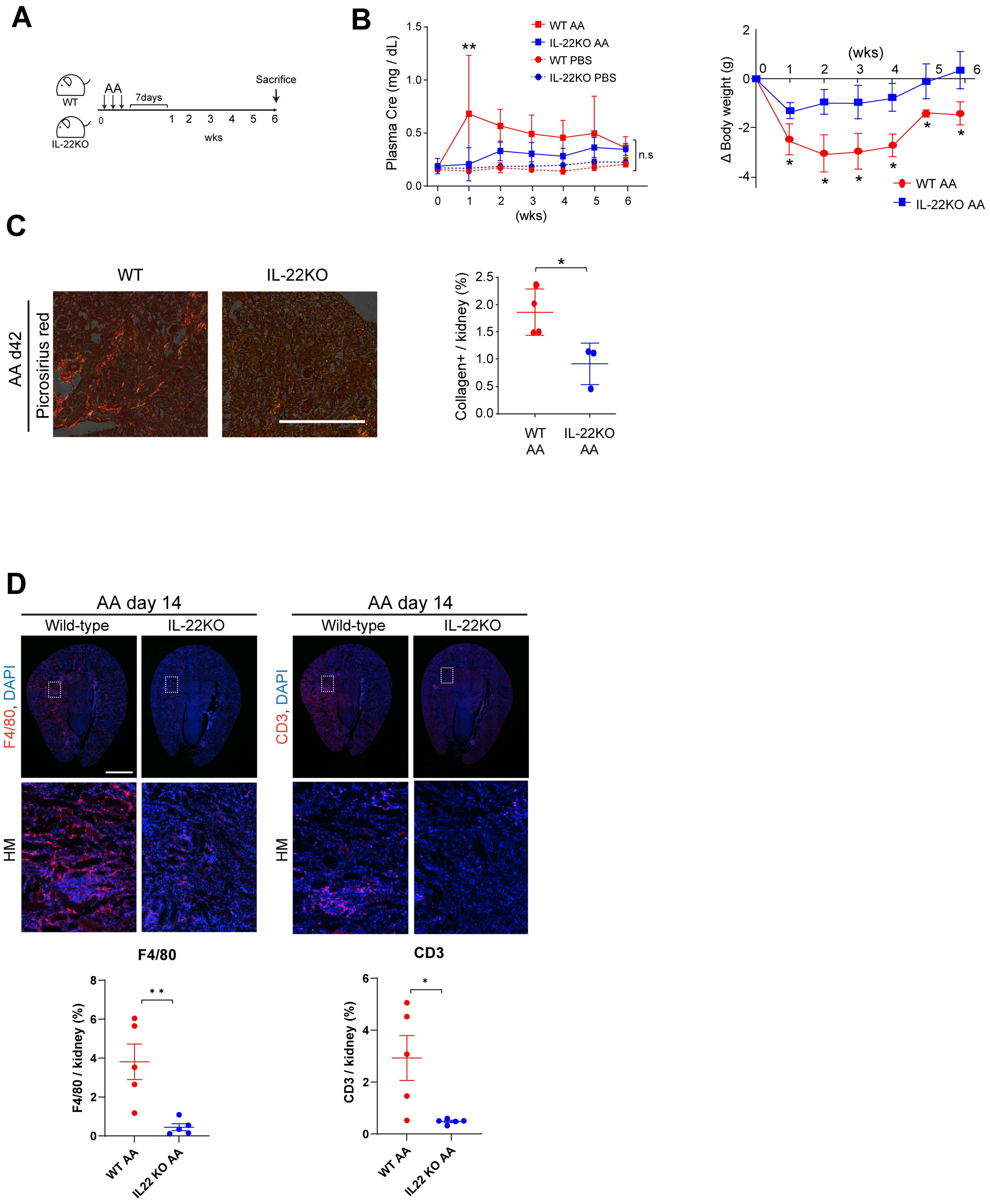

### Supplementary Figure 3

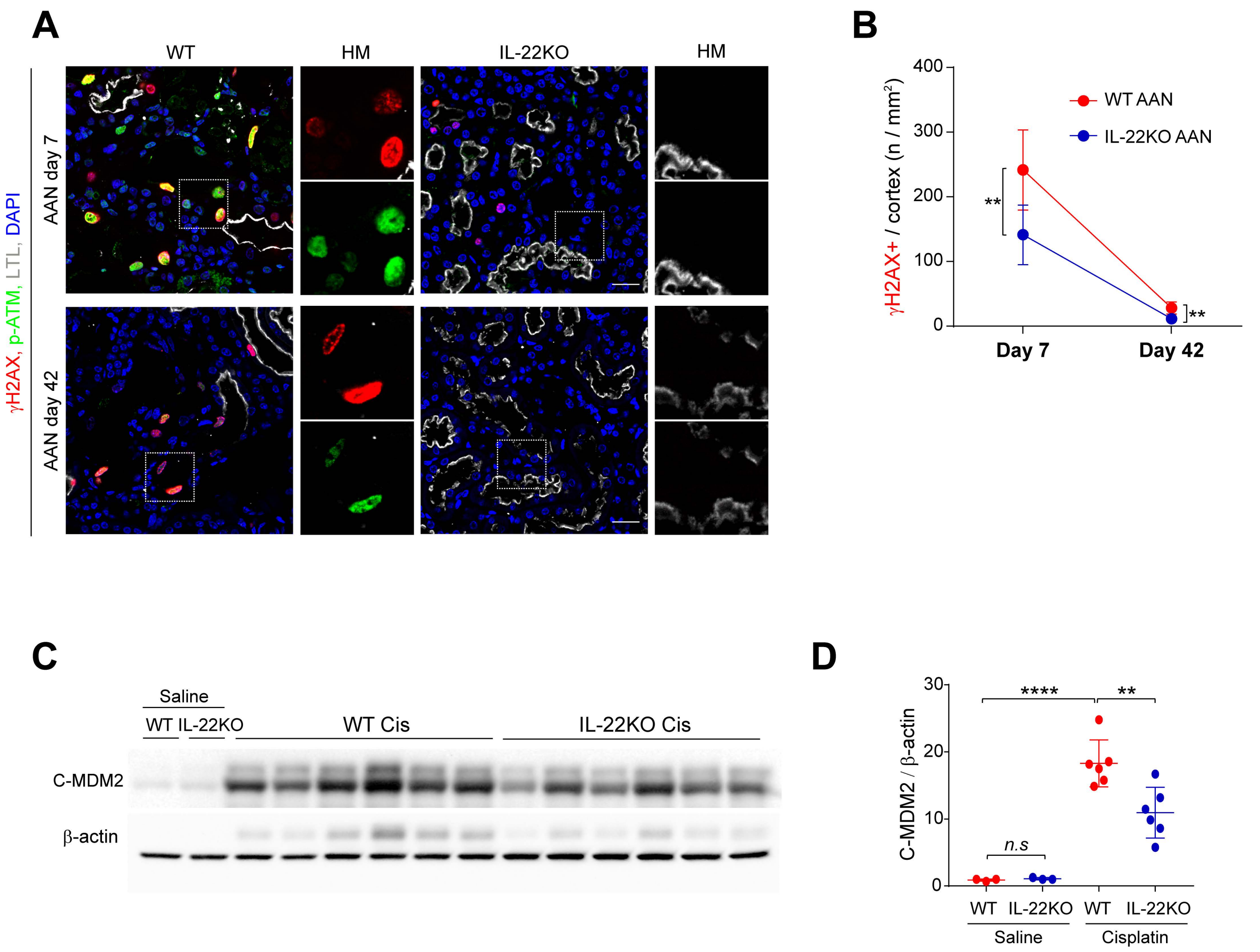

### Supplementary Figure 4

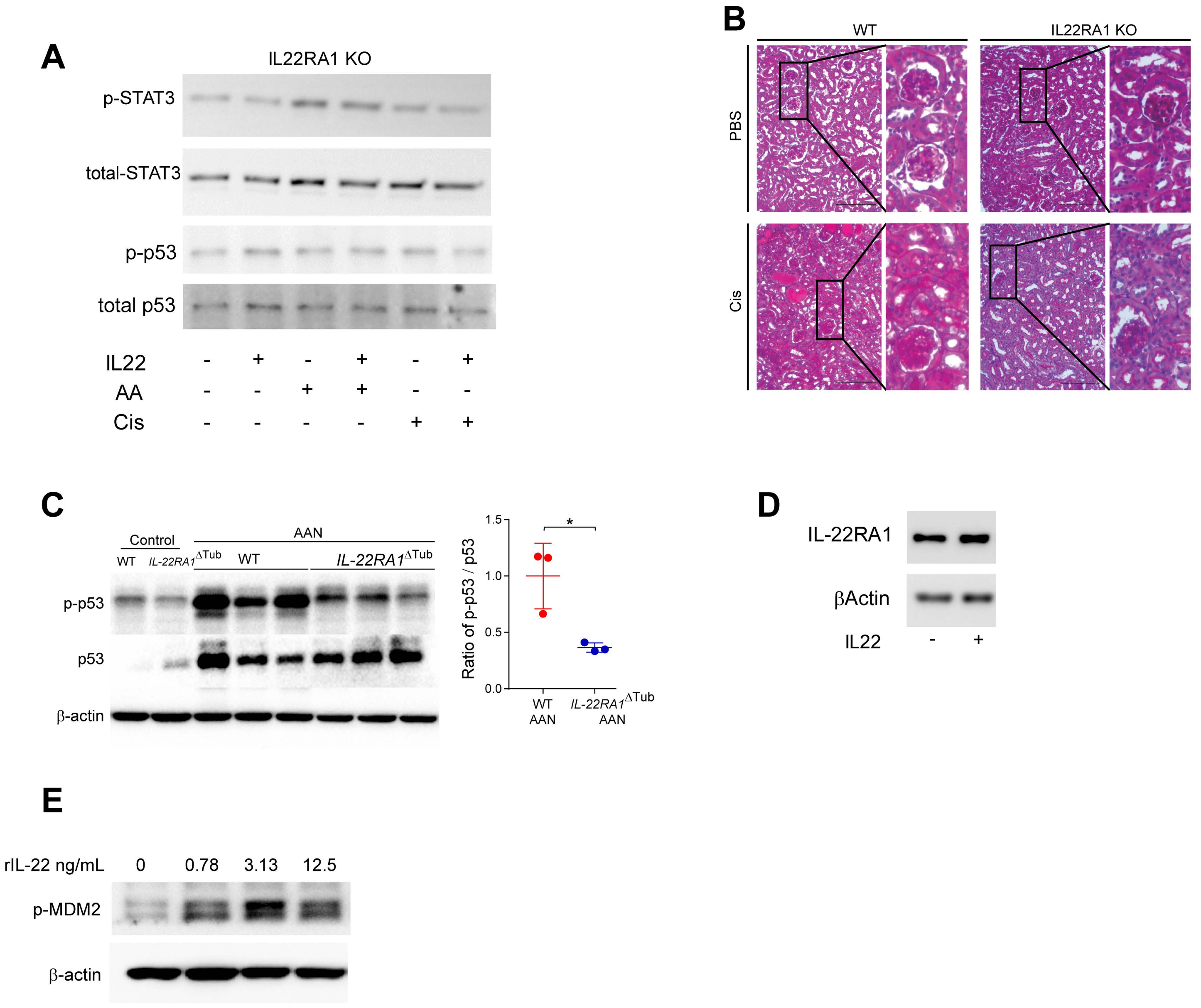

### Supplementary Figure 5

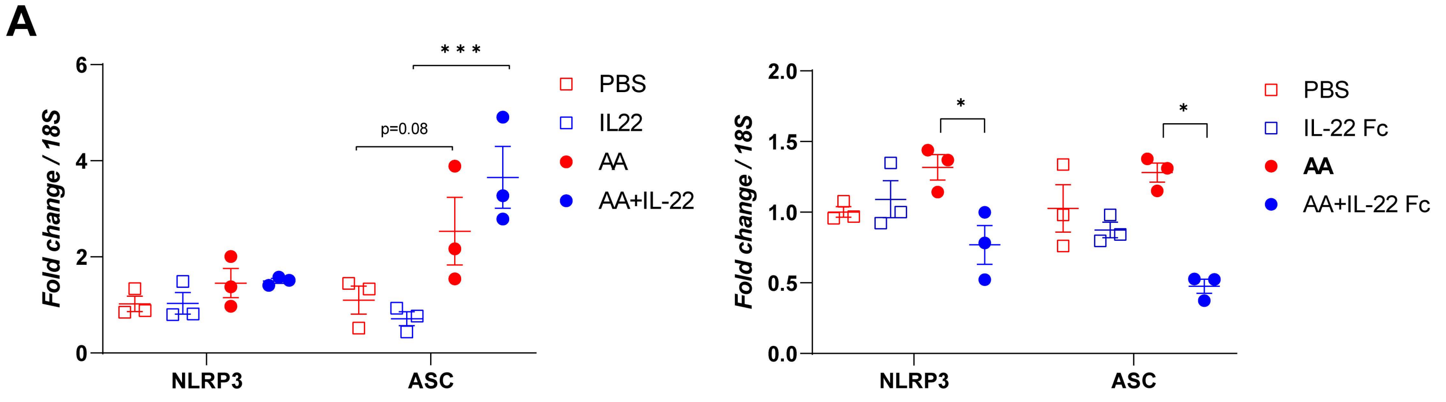


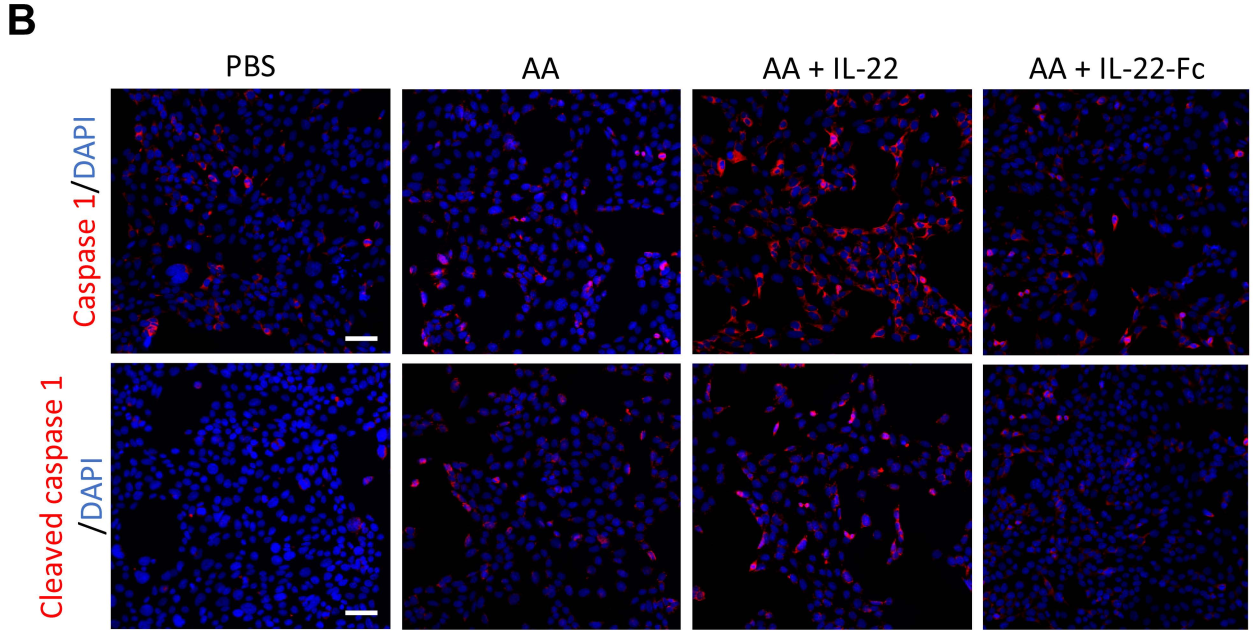


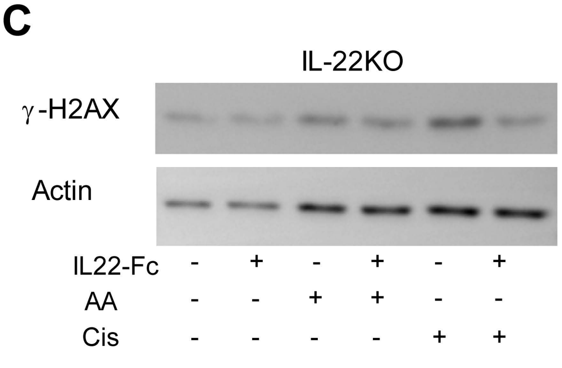

Supplemental Table 1

| Sample ID | Ethnicity | Age | Gender | eGFR (ml/min/1.73m2) | sCr (umol/l) | AKI cause |
|---|---|---|---|---|---|---|
| 20N106162 | Caucasian | 60 | F | 12 | 330 | Cisplatin |
| 20NA06163 | Caucasian | 71 | F | 22 | 231 | Cisplatin |
| 17NA04420 | Caucasian | 55 | F | 17 | 248 | Cisplatin |

Characteristics of patients with cisplatin induced nephropathy. eGFR, estimated glomerular filtration rate;, sCr, serum creatinine; AKI, acute kidney injury

Supplemental Table 2

| Sample ID | Ethnicity | Race | Gender | Cr (mg/dL) | Interstitial fibrosis, tubular atrophy | KIM-1 |
|---|---|---|---|---|---|---|
| UR70 | Unknown / Not Reported | White | Female | 1.65 | 11-25% | + |
| UR76 | NOT Hispanic or Latino | Unknown / Not Reported | Male | 3.76 | > 50% | + |
| UR109 | Unknown / Not Reported | White | Male | 1.04 | 26-50% | + |

Characteristics of patients with or without CKD. CKD, chronic kidney injury; Cr, creatinine; KIM-1, kidney injury molecule-1.

**Supplemental Methods:**

***Renal function.*** Plasma creatinine was measured by Isotope Dilution LC-MS/MS (UAB Biochemical Genetics Laboratory, AL, USA). Plasma BUN was evaluated using QuantiChrom™ Urea Assay Kit (BioAssay Systems, cat# DIUR-100) following the manufacturer's instructions.

***Primary PTC.*** Primary PTC were harvested from wild type and IL-22KO mice as described before [1,2]. Briefly, mice were perfused with sterile PBS and kidney cortex were dissected, minced into small pieces, and incubated with collagenase (1 mg/mL) and trypsin inhibitor (1 mg/mL) in oxygenized media at 37°C with vigorous shaking. After washing with Hank's solution and cultured media, they were centrifuged in 32% Percoll and diluted in DMEM/F12 to separate PTC from other fractions. The isolated PTC were plated on collagen-coated dishes and maintained in primary PTC's media (DMEM/F-12 supplemented with 10%FBS, penicillin/streptomycin, ITS, 0.05 μM hydrocortisone, and 50 μM Vitamin C). Primary PTCs were maintained in DMEM/F-12 supplemented with 2.5% FBS and treated with AA (5-10 mg/mL), cisplatin (0.5 μM), and/or recombinant human IL-22 (rIL-22, Peprotech, cat#; 200-22, NJ) for 2 days up to 7 days. AA was dissolved in sterile PBS at 0.5mg/mL and cisplatin was reconstituted with sterile saline at 1mg/mL. S3I-201 (Millipore-sigma, cat# 573102) was used at 10 mg/mL for inhibition of STAT3 and pifithrin-α (Millipore-sigma, cat# 506132) at 50 μM for p53 inhibition.

***Cell culture.*** RPTEC were maintained in DMEM supplemented with 2.5% FBS and interferon gamma at 33°C. The cells were transferred to 37°C for differentiation and further experiments. RPTEC were incubated with DMEM/F-12 without FBS overnight. Medium was replaced with DMEM/F-12 supplemented with 2.5% FBS and then AA (5 mg/mL), cisplatin (0.1 μM), or rIL-22 were added.

***Real-time PCR.*** Total RNA was isolated from kidney tissue, primary PTC, and cultured cell lines using QIAGEN Mini Kit (Cat #74106) and cDNA was synthesized using iScript™ cDNA Synthesis Kit (BioRad

Cat# 1708891). Real-time PCR was conducted with iTaq Universal SYBR Green (BioRad Cat# 1725121) by Bio-Rad CFX96 Touch Real-Time PCR Detection System with the primers as following; The relative mRNA expressions of each gene were calculated using the ΔΔCt method. Primers used are following; ATM forward, 5'-GCGACTTCAGCTTTCAGGAG-3'; ATM reverse, 5'-TCAACGAGGTGTTTGGTGAG-3'; ATR forward, 5'-CTGTAGCGTCCTTTCGTTCC-3'; ATM reverse, 5'-GCACTTACTCCAGCCACTCC-3'; PUMA forward, 5'-TTCTCCGGAGTGTTCATGC-3'; PUMA reverse, 5'-TACAGCGGAGGGCATCAG-3'; IL-22RA1 forward, 5'-GGACACATCCGGTCTCCTT-3'; IL-22RA1 reverse, 5'-CGGGTCTCCATAGTCAGGTT-3'; IL-22 forward, 5'-CAGACAGGTTCCAGCCCTAC-3'; IL-22 reverse, 5'-CGCCTTGATCTCTCCACTCT-3'; IL-6 forward, 5'-TCGGAGGCTTAATTACACATGTT-3'; IL-6 reverse, 5'-GCATCATCGTTGTTCATACAATC-3'; 18s forward, 5'-GTAACCCGTTGAACCCCATT -3'; 18s reverse, 5'-CCATCCAATCGGTAGTAGCG -3'; GAPDHforward,5'-TCCGCCCCTTCTGCCGATG-3'; GAPDH reverse, 5'-CACGGAAGGCCATGCCAGTGA-3'.

***Western blotting.*** Proteins were isolated with T-PER™ Tissue Protein Extraction Reagent (cat#78510; Thermo fisher scientific) for kidney tissue and M-PER™ Mammalian Protein Extraction Reagent (cat#78501; Thermo fisher scientific) for primary PTC and cultured cell lines. Protease inhibitor cocktail and phosphatase inhibitors were added to the buffers just prior to use. Kidney lysates were centrifuged at 14,000 g for 10 min, and the supernatants were collected. Protein concentrations were determined using Pierce™ BCA protein assay kit (Cat# 23225; Thermo Fisher Scientific). The proteins were separated by SDS-polyacrylamide gel electrophoresis and transferred onto Immobilon-FL polyvinylidene fluoride membranes (Millipore). The membranes were blocked in 5% skim milk / TBS-T or 5% BSA / TBS-T for an hour at room temperature and incubated with antibodies directed against C-casp3 (Cell Signaling Technology Cat# 9664, RRID:AB_2070042), KIM-1 (R and D Systems Cat# AF1817, RRID:AB_2116446), Total p53 (Cell Signaling Technology Cat# 2524, RRID:AB_331743), p-p53 (Ser15) (Cell Signaling Technology Cat# 9286, RRID:AB_331741), MDM2 (Santa Cruz Biotechnology Cat# sc-

965, RRID:AB_627920), p-STAT3 (Ser705) (Cell Signaling Technology Cat# 9145, RRID:AB_2491009), Phospho-Histone H2AX (Ser139) (Cell Signaling Technology Cat# 9718, RRID:AB_2118009), as well as p-ATM (Santa Cruz Biotechnology Cat# sc-47739, RRID:AB_781524) at 4°C overnight. The protein bands were visualized using HRP-conjugated secondary antibodies with Immobilon crescendo western HRP substrate (cat#; WBLUR0500, Millipore, USA). Protein expression levels of β-actin was used as internal control.

***Immunofluorescence staining.*** Paraffin-embedded kidney tissue was sectioned at 4–6 μm thickness and mounted on glass slides. The sections were dried on plate warmer at 48°C overnight. Antigen retrieval was performed using pressure cooker with TE buffer (pH 9.0) or citrate buffer (pH 6.0). The sections were incubated with 1% Triton, 3% Donkey serum in 1% BSA / TBS-T as blocking for an hour at room temperature and then incubated with primary antibodies at 4°C overnight. After washing with PBS, the sections were followed by incubation with secondary antibodies at room temperature for an hour. Antibodies directed against KIM-1 (R and D Systems Cat# AF1817, RRID:AB_2116446), C-casp3 (Cell Signaling Technology Cat# 9664, RRID:AB_2070042), and p-ATM (Santa Cruz Biotechnology Cat# sc-47739, RRID:AB_781524) were used as primary antibodies. Cy3-Donkey anti-mouse IgG antibody (Jackson ImmunoResearch Labs Cat# 715-166-151, RRID:AB_2340817), DyLight 488-donkey anti mouse IgG antibody (Jackson ImmunoResearch Labs Cat# 715-486-151, RRID:AB_2572300), Cy3-donkey anti-mouse IgG antibody (Jackson ImmunoResearch Labs Cat# 711-166-152, RRID:AB_2313568), DyLight 488-donkey anti rabbit IgG antibody (Jackson ImmunoResearch Labs Cat# 711-485-152, RRID:AB_2492289), and Alexa Fluor 647-donkey anti-goat IgG antibody (Jackson ImmunoResearch Labs Cat# 705-606-147, RRID:AB_2340438) were used as secondary antibodies. Stained whole kidneys were scanned by NIKON eclipse Ti2-E automated microscope. In addition, images were also acquired using Zeiss 710 confocal microscope. For each sample, targeted protein-positive area or the number of targeted protein-positive cells were analyzed by NIS-Elements Microscope Imaging Software and divided by cortex or kidney area to obtain proportion of positive area (%) or positive cell number (n /$mm^2$).

***PAS staining***. Paraffin-embedded kidney tissue was sectioned at 4–6 μm thickness and mounted on glass slides. The sections were dried on plate warmer at 48°C overnight. Sections were deparaffinized and rehydrated with Histo-Clear (cat#; HS-200, National Diagnostics, GA) or xylene and graded ethanol series, respectively. The sections were then immersed in a 1% periodic acid solution (cat#; PAQ250, ScyTek Laboratories, UT) for 10 minutes at room temperature, followed by rinsing in distilled water. Next, the sections were placed in Schiff's reagent for 30 minutes at room temperature, rinsed again in hot tap water for 5 minutes, and counterstained with Mayer's hematoxylin for 1 minutes. Finally, the sections were dehydrated in graded ethanol series, cleared in Histo-Clear or xylene, and mounted with a coverslip.

***Picrosirius red staining.*** Paraffin-embedded kidney tissue was sectioned at 4–6 μm thickness and mounted on glass slides. The sections were dried on plate warmer at 48°C overnight. After de-paraffinization, the sections were incubated with Weigert's hematoxylin for 8 minutes to stain nuclei, followed by incubation with Sirius red solution (cat#; 26357-02, electron microscopy science, PA) for three hours to stain collagen fibers. Stained sections were imaged with NIKON eclipse Ti2 equipped with a linear polarized lens and collagen deposition area / kidney was analyzed by NIS-Elements Microscope Imaging Software.

***Counting p-ATM+ foci in vitro.*** The number of green p-ATM foci in nucleus were scored manually under a Nikon Eclipse Ti2-E automated microscope at x40. For each sample, approximately 20 nuclei were analyzed and the average calculated.

***Urine IL-22.*** IL-22 concentration was measured by ELISA (cat#; M2200, R&D systems, MN). Kidney tissue IL-22 samples were prepared from kidney lysates at the same concentration with BCA protein assay (cat#; 23225, Thermo Fisher scientific).
